# Sex-specific evolution of a *Drosophila* sensory system via interacting *cis-* and *trans-*regulatory changes

**DOI:** 10.1101/2022.01.12.475924

**Authors:** David Luecke, Gavin Rice, Artyom Kopp

## Abstract

The evolution of gene expression via cis-regulatory changes is well established as a major driver of phenotypic evolution. However, relatively little is known about the influence of enhancer architecture and intergenic interactions on regulatory evolution. We address this question by examining chemosensory system evolution in *Drosophila*. *D. prolongata* males show a massively increased number of chemosensory bristles compared to females and males of sibling species. This increase is driven by sex-specific transformation of ancestrally mechanosensory organs. Consistent with this phenotype, the *Pox neuro* transcription factor (*Poxn*), which specifies chemosensory bristle identity, shows expanded expression in *D. prolongata* males. *Poxn* expression is controlled by non-additive interactions among widely dispersed enhancers. Although some *D. prolongata Poxn* enhancers show increased activity, the additive component of this increase is slight, suggesting most changes in *Poxn* expression are due to epistatic interactions between *Poxn* enhancers and trans-regulatory factors. Indeed, the expansion of *D. prolongata Poxn* enhancer activity is only observed in cells that express *doublesex* (*dsx*), the gene that controls sexual differentiation in *Drosophila* and also shows increased expression in *D. prolongata* males due to *cis*-regulatory changes. Although expanded *dsx* expression may contribute to increased activity of *D. prolongata Poxn* enhancers, this interaction is not sufficient to explain the full expansion of *Poxn* expression, suggesting that cis-trans interactions between *Poxn*, *dsx*, and additional unknown genes are necessary to produce the derived *D. prolongata* phenotype. Overall, our results demonstrate the importance of epistatic gene interactions for evolution, particularly when pivotal genes have complex regulatory architecture.

**Research Highlights:** In *Drosophila prolongata* males, many mechanosensory organs are transformed into chemosensory. This is due in part to interacting regulatory changes in *Poxn*, which controls chemosensory organ development, and *dsx*, which controls sexual differentiation.

**Graphical Abstract:** 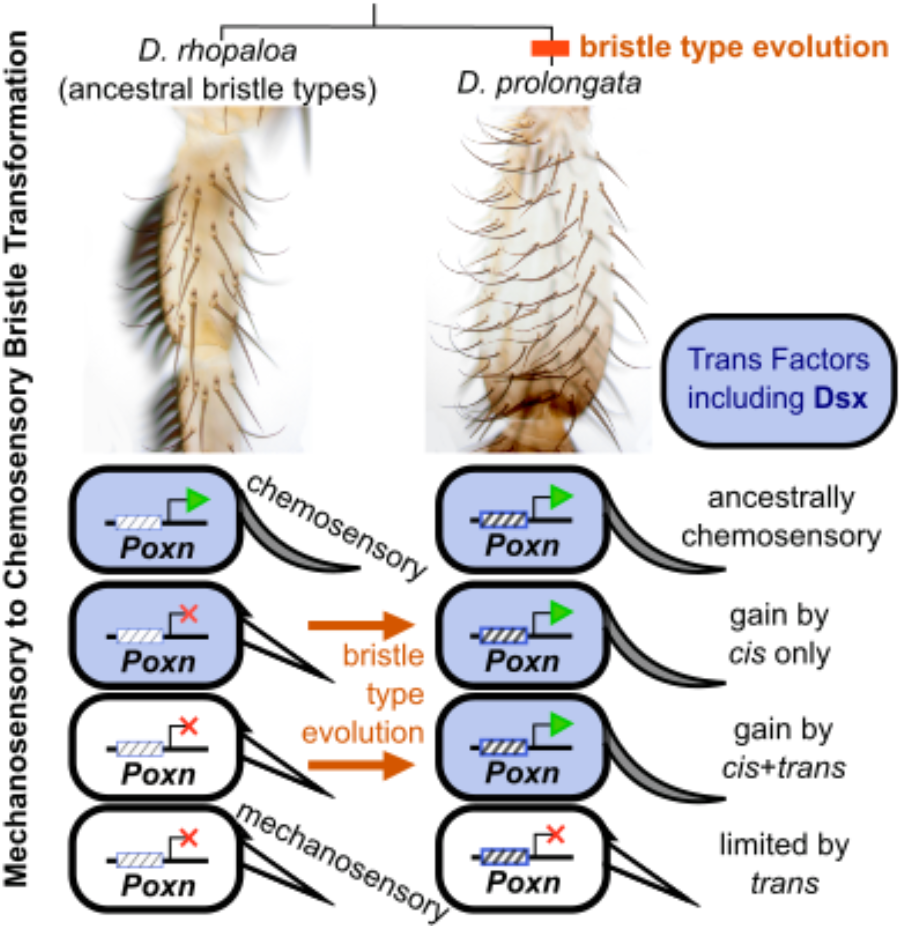

## Introduction

During the evolution of biological diversity novel morphologies often result from redeployment of conserved developmental modules into new spatial and temporal contexts (Monteiro & Podlaha, 2009). Changes in the expression of “master regulatory genes”, *i.e.* genes that initiate the expression of downstream gene networks, play an especially important role in this model of morphological evolution (Fisher et al., 2019; Gao & Davidson, 2008; Glassford et al., 2015; Niwa et al., 2010; Oliver et al., 2012; Vlad et al., 2014). Many, if not most, evolutionary changes in the expression of master regulatory genes are due to mutations is the *cis-*regulatory elements (CREs, or enhancers) of these genes (Carroll, 2008; Monteiro & Gupta, 2016). Although new gene expression domains most commonly evolve via modification of existing enhancers, the origin of new CREs can also contribute to this process (Rebeiz et al., 2011; Rebeiz & Tsiantis, 2017).

An important and largely unexplored question is how a gene’s preexisting landscape of CREs influences the paths and outcomes of regulatory evolution (Gao & Davidson, 2008; Sabarís et al., 2019). In this report, we address this question by examining the male-specific expansion of chemosensory system in the fruit fly *Drosophila prolongata*. In *D. prolongata* males, the number of chemosensory bristles on the prothoracic legs is far greater than in females, or in males of any other *Drosophila* species including its closest relatives (Figure 1). The spatial arrangement of these bristles is also unusual. In other *Drosophila* species, mechanosensory bristles are arranged in longitudinal rows with single chemosensory bristles interspersed between these rows, reflecting the ancestral *Drosophila* condition (Held, 1979). However, in male *D. prolongata* several longitudinal rows are composed of both mechanosensory and chemosensory bristles. This pattern can best be explained by redeployment of the chemosensory bristle development program into new locations, including ancestrally mechanosensory bristles. The trait’s sex-limited nature suggests novel regulatory interactions with the sexual differentiation pathway (Hopkins & Kopp, 2021). These changes are part of a larger suite of male-specific modifications of *D. prolongata* forelegs, which also include greatly increased size and unusual pigmentation (Luecke & Kopp, 2019). The recency of these changes, together with existing knowledge of bristle development and sexual dimorphism in *Drosophila*, make the chemosensory system of *D. prolongata* a promising model for investigating the evolutionary redeployment of developmental modules.

**Figure 1:**
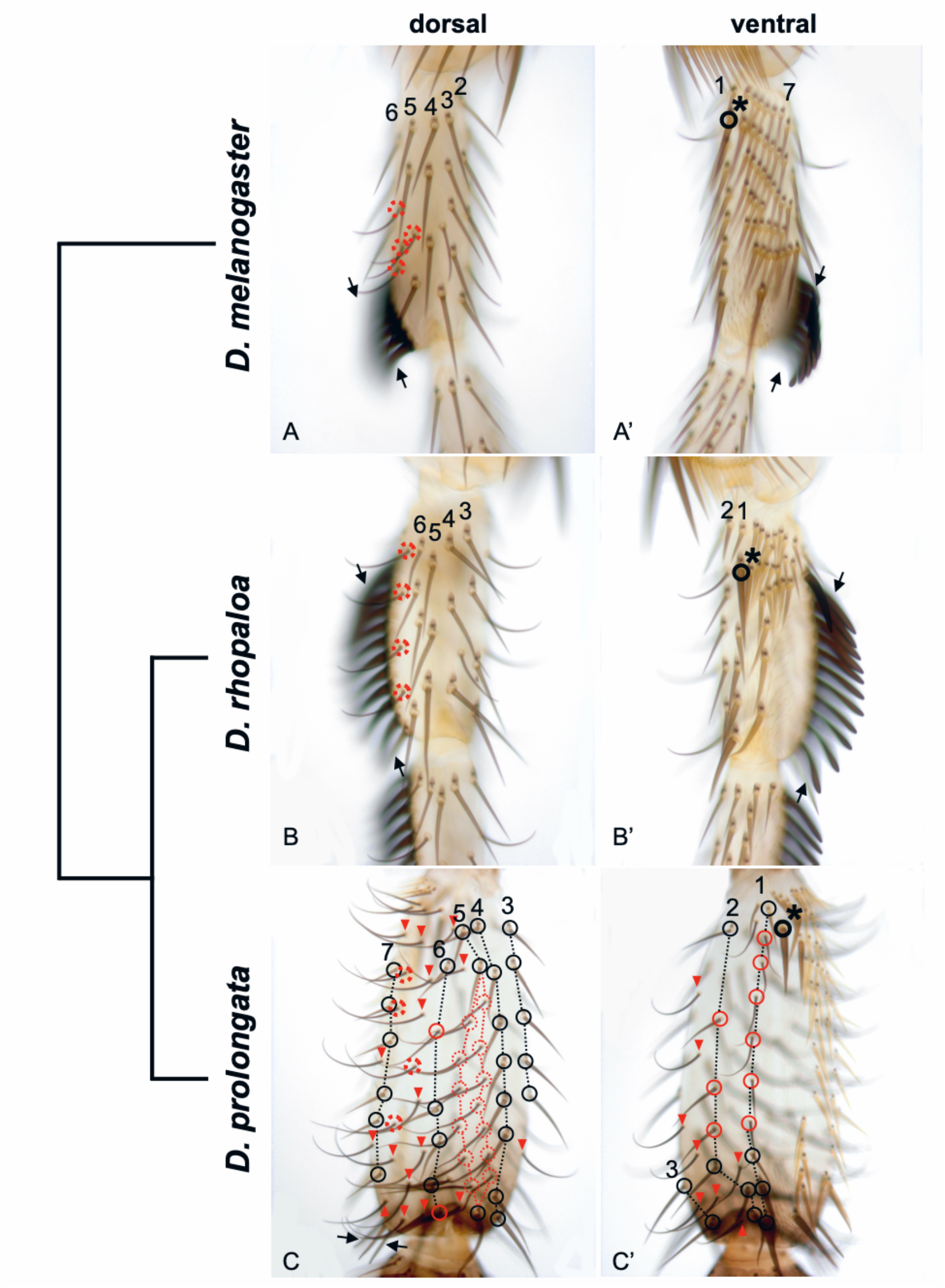
Recent male-specific expansion of the leg chemosensory system in *D. prolongata*. Images of the first tarsal segment (t1) of the male prothoracic leg of three species, with a cladogram indicating their evolutionary relationships. The male foreleg in *D. prolongata* has a derived size increase, pigmentation pattern, and bristle organization, all apparent in these images; this investigation focuses on derived patterning of bristles. Two landmarks are clearly homologous across species: the conserved 1’ bristle (Tokunaga, 1962) in proximal t1 (thick black circle with asterisk), and the most distal transverse bristle row (TBR), which is modified into a sex comb in *D. rhopaloa* and *D. melanogaster* but not in *D. prolongata* (arrows). Homologous longitudinal bristle rows 1-7 (Tokunaga, 1962) can be identified relative to these landmarks despite their morphological divergence. Mechanosensory bristles, circled in black, have basal bracts and are straight and pointed. Chemosensory bristles, circled in red, are distinguished by their curved shape and the absence of bracts. A, A’. *D. melanogaster.* All 7 longitudinal rows are made up exclusively of mechanosensory bristles. The anterior-ventral side between longitudinal rows 1 and 7 carries a series of TBRs, the most distal of which develops into a rotated sex comb (Kopp, 2011; Tokunaga, 1962). A small number of chemosensory bristles are present on the dorsal leg surface; sex-specific chemosensory bristles (Mellert et al., 2012; G. R. Rice et al., 2019; Tokunaga, 1962) are circled in red dashed lines. B, B’. *D. rhopaloa.* Longitudinal rows 1-6 are composed of only mechanosensory bristles. Most TBRs develop into a large rotated sex comb (Tanaka et al., 2009). Sex comb rotation displaces longitudinal row 7 as well as dorsal chemosensory bristles. The chemosensory bristles circled in red dashed lines are homologous to the ones circled in *D. melanogaster* (A). C, C’. *D. prolongata*. Bristles that belong to identifiable longitudinal rows are circled and connected with dotted lines. Rows 1, 2, 5, and 6 include both mechanosensory (black) and chemosensory (red) bristles. Row 5 runs along two parallel rows of chemosensory bristles, either of which could be homologous to row 5 in other species (the other being supernumerary); this ambiguity is depicted as dotted red circles and lines. Additional chemosensory bristles between longitudinal rows 6 and 7 (circled in red dashed lines) are probably homologous to male-specific chemosensory bristles in *D. melanogaster* and *D. rhopaloa* (red dashed circles in A, B). Only vestigial TBRs remain at the distal (black arrows) and proximal (near the 1’ bristle) ends of t1. Many supernumerary chemosensory bristles are interspersed between the conserved longitudinal rows, forming additional partial rows (red arrowheads and one of the row 5 branches).

In *Drosophila* adults the most abundant external sensory organs are mechanosensory bristles, which are innervated by a single neuron and have straight and pointed shafts with a basal bract. A smaller set of bristles on the legs, wings, and genitalia are chemosensory; these are polyinnervated, and can be identified by curved shafts and the lack of bracts (Nayak & Singh, 1983). Bristle specification begins with the establishment of proneural clusters (PNCs) – groups of cells from which a single sensory organ precursor (SOP) emerges (Furman & Bukharina, 2008; Simpson, 1990). The SOP fate is then maintained by a feedback loop involving the transcription factor *senseless* (Jafar-Nejad et al., 2003; Nolo et al., 2000). Chemosensory bristles are distinguished from mechanosensory by expression of the transcription factor *Paired-box neuro* (*Pox neuro* or *Poxn*) (Bopp et al., 1989; Dambly-Chaudière et al., 1992). Ectopic *Poxn* expression converts mechanosensory SOPs to chemosensory bristle fate, and can induce supernumerary chemosensory bristles, whereas loss of *Poxn* causes chemosensory bristles to be either re-specified as mechanosensory or lost altogether (Awasaki & Kimura, 1997; Nottebohm et al., 1992, 1994). Thus, *Poxn* is necessary and sufficient for chemosensory fate within SOPs, and may also contribute to maintaining SOP commitment.

In the prothoracic leg of the model species *D. melanogaster*, the mechanosensory bristle pattern consists of seven longitudinal rows on the anterior, dorsal, and posterior sides, and a series of transverse bristle rows (TBRs), used in grooming, on the ventral side (Hannah-Alava, 1958; Shroff et al., 2007). In males, the most distal TBR is modified into a thick, heavily pigmented, and rotated sex comb (Held, 2002). Chemosensory bristles are interspersed between the rows in a stereotypic but less organized pattern, with a greater number in males than females (Hannah-Alava, 1958; Nayak & Singh, 1983). Although the number of bristles varies across species, the overall spatial pattern is highly conserved (Held, 1979). Interspecific variation in the *melanogaster* species group, which includes *D. prolongata*, is dominated by differences in the position and size of the sex comb (Kopp, 2011; Kopp & True, 2002). Chemosensory leg bristles also vary in number and position across species, usually more numerous in males (Kopp and Barmina, unpublished). The leg bristle pattern is established early in pupal development, with chemosensory bristles specified earlier than all but the largest mechanosensory ones (Held, 1990; Rodríguez et al., 1990). In the developing leg, PNCs are first seen as isolated clusters before coalescing into stripes along the proximal-distal axis (Orenic et al., 1993), yielding the typical *Drosophila* pattern – rows of mechanosensory bristles interleaving scattered chemosensory bristles.

Male and female bristle pattern variation is controlled by the *doublesex* (*dsx*) transcription factor, responsible for most morphological sexual dimorphism in *Drosophila* and other insects (Hopkins & Kopp, 2021; Robinett et al., 2010). Variation in *dsx* expression controls differences in the presence and morphology of the sex comb across *Drosophila* species (Kopp, 2011; Tanaka et al., 2011). Importantly, *dsx* is also responsible for sexual dimorphism in chemosensory bristle number, and specifically the development of extra male-specific chemosensory bristles absent in females (Mellert et al., 2012).

This background suggests the male-specific increase in the number of chemosensory bristles in *D. prolongata* forelegs involves an expansion of *Poxn* expression into leg regions that ancestrally produced mechanosensory bristles, and that this expansion is controlled at least partly by *dsx*. *dsx* is expressed in the foreleg in a tightly restricted spatial pattern (Robinett et al., 2010; Tanaka et al., 2011) specified by at least three CREs (Rice et al., 2019). An early enhancer drives expression in all sexually dimorphic leg tissues, including the sex comb and sex-specific chemosensory bristles, during the late larval and early pupal stages when SOP specification occurs. Later in pupal development *dsx* expression is controlled by two separate CREs – the late sex comb enhancer and the chemosensory bristle enhancer (Rice et al., 2019).

Regulation of *Poxn,* which has many functions in development beyond chemosensory organs (Awasaki & Kimura, 1997, 2001; Boll & Noll, 2002; Glassford et al., 2015), is even more complex. Using a series of transgenic rescue constructs (Boll & Noll, 2002) showed that temporally distinct early and late phases of *Poxn* expression in the leg are driven by different enhancers, with early expression controlling bristle specification. They found that the genomic region responsible for early expression is a composite of at least three CREs spread over more than 5kb of noncoding DNA (Figure 3a). The “core bristle” region identified by (Boll & Noll, 2002) is required for bristle development, but this region alone was not sufficient to produce the complete set of chemosensory bristles in their experiments. Full chemosensory bristle development required the addition of a separate region we label “over-rescue,” as the addition of this region was necessary for *Poxn* expression in the full wild-type pattern, but also generated ectopic expression beyond the normal domain. Repression of this ectopic expression requires additional intronic and downstream sequences (Boll & Noll, 2002). In short, *Poxn* expression in the leg cannot be “disassembled” into additive modules. Rather, the minimal functional element for correct *Poxn* expression in leg bristles appears to be the entire *Poxn* locus, or at least multiple interacting regions distributed throughout.

**Figure 3:**
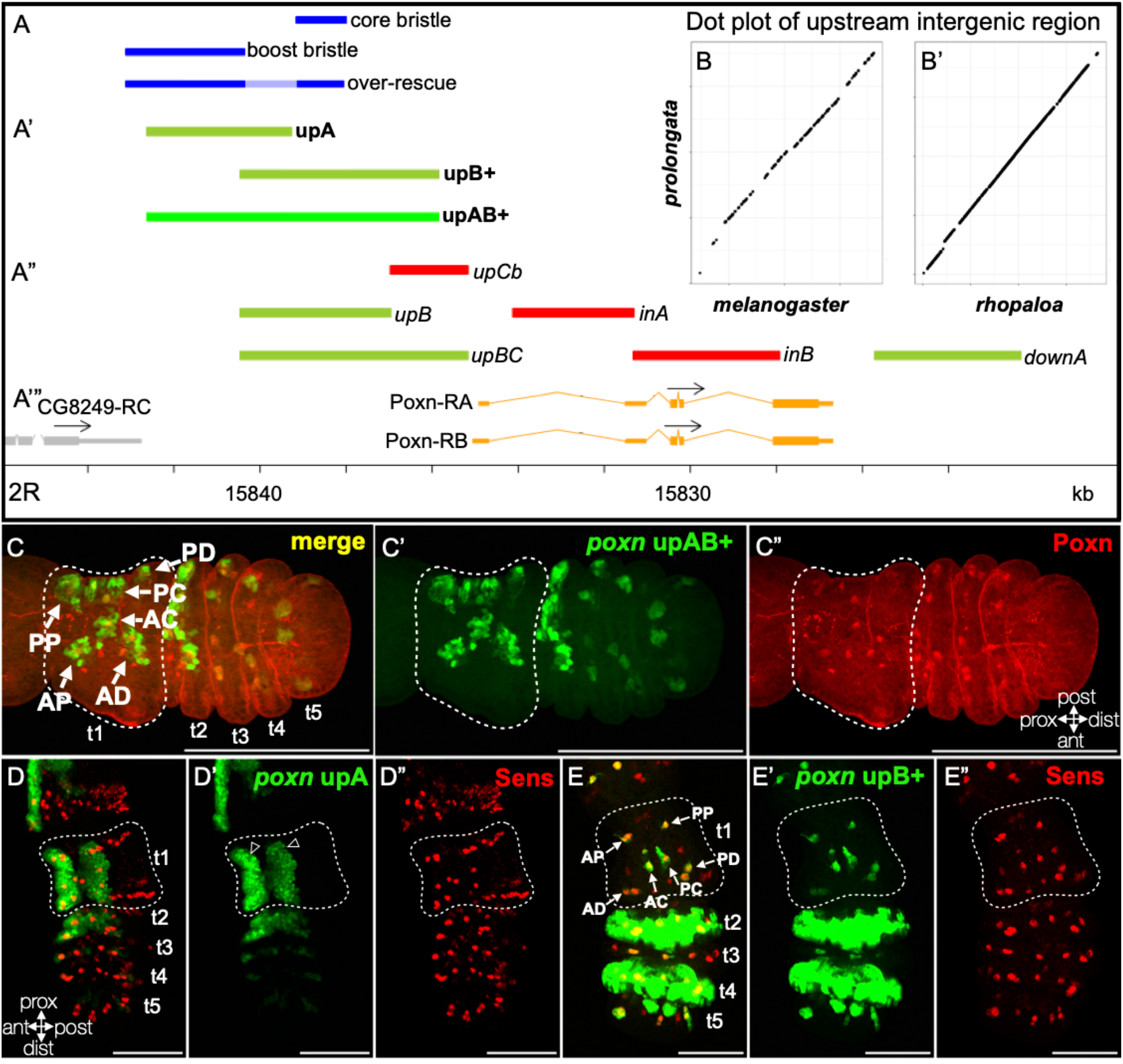
Complex *cis*-regulatory organization of *Poxn* in *D. melanogaster*. The *Poxn* locus, including upstream and downstream non-coding regions, is shown in A’’’, with the *Poxn* transcripts in orange. Reporter constructs (A-A’’) are aligned with the locus schematic. A. Previously described regulatory regions in *D. melanogaster* (Boll & Noll, 2002) reflect the complex regulation of *Poxn* in leg chemosensory bristles. In rescue constructs consisting of *Poxn* CDS under the control of *Poxn* non-coding regions in a *Poxn* mutant background, the “core bristle” region is required to recover any chemosensory bristles, but it is not alone sufficient to produce the full wild-type complement of bristles. The “boost bristle” region together with the “core bristle” element (the “over-rescue” construct) rescues the full set of chemosensory bristles, but also results in ectopic bristles. The light blue region of the “over-rescue” fragment can be deleted with no effect on *Poxn* leg bristle phenotype (Boll & Noll, 2002). A’. Key reporter fragments examined in this study (bold labels). A’’ Additional reporter fragments described in Figure S3 (italic labels). Reporters that drive expression in pupal legs are in green, and regions that lack leg expression are in red. B, B’. The intergenic region between CG8249 and *Poxn* is syntenic in *D. melanogaster*, *D. rhopaloa*, and *D. prolongata*. No large inversions, insertions, or deletions have occurred in this region, indicating that upstream regulatory elements are homologous between species. C. Z-stack of the dorsal male T1 leg surface at 5 hr APF showing the activity of the *D. melanogaster* upAB+ fragment in *D. melanogaster*. Anterior is down, t1 segment is outlined. UAS-GFP.nls driven by the *Poxn*-Gal4 reporter is in green, and Poxn protein expression in red. The *Poxn* upAB+ enhancer produces six clusters of expression in t1, each marked with an arrow and labeled based on its position: anterior-proximal - AP, anterior-distal - AD, anterior-central - AC, posterior-central - PC, posterior-proximal - PP, and posterior-distal - PD. These clusters correspond to Poxn*-*expressing SOPs (C’’, guinea pig Poxn), but extend beyond the SOP itself (C, C’). Several anterior Poxn-expressing SOPs lie outside of these expression clusters. D. Z-stack of the dorsal T1 male leg surface at 5 hr APF showing the activity of the *D. melanogaster* upA region in *D. melanogaster*. *Poxn*-Gal4 > UAS-GFP.nls is in green, and Sens in red. Reporter expression is seen in two bands of epithelial cells along the dorsal-anterior and dorsal-posterior surfaces of t1 and t2. Expression is stronger in epithelial cells, and weaker in the SOPs marked by Sens (see holes of weak expression corresponding to Sens-expressing cells, marked with outlined arrowheads). E. Z-stack of the dorsal T1 male leg surface at 5 hr APF showing the activity of the *D. melanogaster* upB+ region in *D. melanogaster*. *Poxn*-Gal4 > UAS-GFP.nls is in green, and Sens in red. Expression is seen in six SOP cells in t1, corresponding to the six clusters generated by the upAB+ enhancer (C’), but without the surrounding epithelial expression. However, upB+ drives ectopic epithelial expression in the more distal tarsal segments.

It is unclear how a complex, non-modular regulatory architecture can enable the evolution of gene expression in isolated developmental contexts (Sabarís et al., 2019). The expansion of chemosensory bristles in *D. prolongata* presents an excellent model to study this question. This recently evolved phenotype may reflect a redeployment of the developmental network controlled by *Poxn*, a gene with a non-modular regulatory landscape, interacting with the sex-differentiation input mediated by *dsx*, with more classically modular regulatory regions. How the difference in preexisting regulatory substrates affects recruitment of these genes into the novel chemosensory bristle domain is an open question. We compare the *Poxn* and *dsx* expression patterns in *D. prolongata*, its close relative *D. rhopaloa*, and *D. melanogaster*, and identify the regulatory regions responsible for species- and sex-specific expression. We provide initial evidence that intergenic interactions have contributed to the origin of the *D. prolongata* phenotype and propose a model for gene expression changes that produced the recruitment of *Poxn* into a new tissue. Our findings suggest that the complex *cis*-regulatory architecture of *Poxn* may act as a constraint on its evolution, such that spatially restricted evolutionary changes in its expression require contributions from other loci.

## Materials and Methods

### Fly stocks

The isofemale strains of *D. prolongata* SaPa001 (collected in SaPa, Vietnam, September 2004), *D. rhopaloa* BaVi067 (BaVi, Vietnam, March 2005), and *D. melanogaster* WI89 (Winters, CA, 1999) were used as wild types. Standard fly strains were obtained from Bloomington Stock Center including UAS-GFP.nls (Bloomington stock #4775), UAS-svRNAi (Bloomington stock #27269), UAS-Dcr2 (Bloomington stock #24650), UAS-Poxn (Jiao et al., 2001), and UAS-Dsx^M^ (Bloomington stock #44224). A double-balanced UAS-Dcr2 / CyO; UAS-svRNAi / MKRS stock was created and used for adult bristle reporter experiments. Transgenic reporter flies were generated by BestGene Inc. using attP2 (Bloomington stock #8622) and attP40 (y^1^ w^67c23^; P{CaryP}attP40) (Markstein et al., 2008) integration sites crossed to germline-driven PhiC31 (Bloomington stock # 40161).

### Antibody production and immunohistochemistry

Polyclonal antibodies against the *D. melanogaster* Poxn protein were produced using the same epitope as in (Bopp et al., 1989). The cDNA sequence encoding this epitope was cloned into the pMAL-p2X vector (a gift from Paul Riggs, Addgene plasmid # 75287), expressed in BL21 cells with IPTG induction and affinity-purified on an amylose column using standard protocols (NEB #E8200S). Guinea pig reactive serum was produced by UC Davis Comparative Pathology Laboratory, and used at 1:10 dilution in combination with mouse anti-Cut (DSHB Hybridoma Product 2B10 deposited by Rubin, G.M.) (Blochlinger et al., 1990) at 1:10 in Antibody Dilution Buffer (1x PBS, 1% BSA, 0.3% Triton X-100). The secondary antibodies were Cy3-anti guinea pig (Sigma-Aldrich) and Dylight 649-anti mouse (Thermo Fisher), both at 1:200 in the same buffer. Stained tissues were treated with Image-iT FX Signal Enhancer (Invitrogen) and washed in 5% normal goat serum in PBS + 0.1% Tween-20. Other primary antibodies used were rabbit anti-Poxn (shared by M. Noll) (Bopp et al., 1989) at 1:50, rat anti-DsxM (shared by B. Oliver) (Hempel & Oliver, 2007) at 1:50, mouse anti-Dsx[DBD] (DSHB Hybridoma Product DsxDBD deposited by Baker, B.S.) (Mellert et al., 2012) at 1:10, and guinea pig anti-Sens at 1:1000 (Shared by H. Bellen) (Nolo et al., 2000), following the protocols described in (Tanaka et al., 2009). Morphological marker staging and leg dissection were done as in (Tanaka et al., 2011) and see stage description in Results.

### Transgenic reporter constructs

The boundaries of reporter fragments were designed to include known regulatory regions (Boll & Noll, 2002; G. R. Rice et al., 2019), and to overlap when possible. Evo-printer (Odenwald et al., 2005) and VISTA (Frazer et al., 2004) were used to design primer sequences with minimal sequence divergence between *D. melanogaster* and *D. rhopaloa*, in an effort to identify homologous enhancer boundaries, based on sequences available on FlyBase (Larkin et al., 2021). Primer sequences are provided in Supplementary Table 1. Synteny dotplots were made with Geneious using *D. prolongata* sequence from a draft genome assembly generated by Dovetail Genomics. Candidate *cis*-regulatory regions were amplified by PCR with OneTaq polymerase (NEB) and cloned into the pCR8 plasmid (Invitrogen), then transferred using the Gateway system into the pBPGUw (Pfeiffer et al., 2008) or pGreenfriend (Miller et al., 2014) vectors for Gal4 or GFP reporters, respectively. Multiple independent clones were produced for each region, and clone boundaries were confirmed with Sanger sequencing from M13 and T7 primers. Reporter vectors were injected by BestGene into either attP2 or attP40; all allele comparisons were done between the same integration site, using two independently cloned reporters.

### Confocal microscopy and image processing

At least ten legs were examined for each enhancer clone, and at least three image stacks were produced for each region; representative images for figures were chosen based on tissue orientation and staining clarity. Confocal images were taken using an Olympus FV1000 laser scanning confocal microscope and processed in ImageJ (Schneider et al., 2012).

### Light microscopy

Legs were dissected from adult flies and mounted in PVA-Mounting-Medium (BioQuip) to clear the samples. Phenotypic effects of sv RNAi were quantified by inspection of slides blinded to enhancer allele under light microscope, with each first tarsal bristle given a score as intact=1, partial knockout=0.5, or fully removed=0; see the Figure 4 legend for sv-RNAi sample sizes.

**Figure 4:**
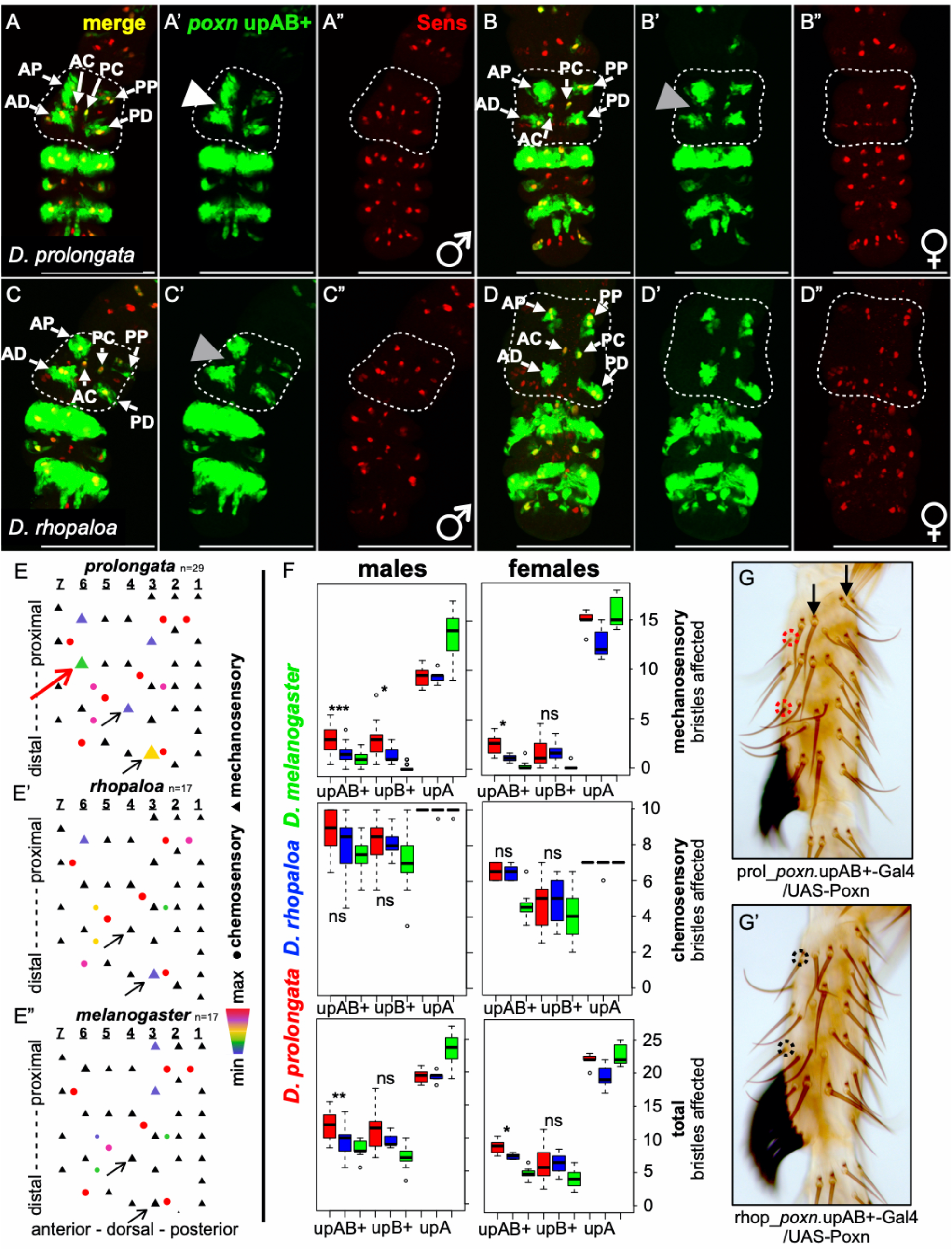
Expanded spatial activity of the *D. prolongata Poxn* leg enhancer. A-D. Z-stacks of male (A, C) and female (B, D) dorsal T1 legs at 5hr APF. All images show transgenic *D. melanogaster* carrying *D. prolongata* or *D. rhopaloa Poxn* reporters. Anterior is to the left, t1 segment is outlined. *Poxn*_upAB+-Gal4/UAS-GFP.nls is in green and Sens (marking SOPs) is in red. upAB+ enhancer activity in t1 is generally similar between the two species and in both sexes; the AP, AD, PP, and PD clusters are similar to the *D. melanogaster* enhancer (Fig. 3 C), while the AC and PC clusters are reduced to isolated cells. In males, expression from the *prolongata* upAB+ enhancer extends into the gap between the AP and AD clusters (white arrowhead in A’); this expansion is not seen in females carrying the *D. prolongata* enhancer, or in males and females carrying the *D. rhopaloa* enhancer (grey arrowheads in B’, C’). *D.* Diagrams showing the effect of UAS-svRNAi expression driven by *Poxn* upAB+ alleles from *D. prolongata* (E), *D. rhopaloa* (E’), and *D. melanogaster* (E”) on the *D. melanogaster* male prothoracic t1 bristle pattern. Mechanosensory bristles are shown as triangles, and chemosensory bristles as circles. Each longitudinal row is numbered as in Fig. 1. Color and size reflect enhancer activity in each bristle, as indicated by the fraction of these bristles impacted in *Poxn*-Gal4/UAS-*sv*RNAi males. The red arrow indicates the region in the mid-anterior tarsus that shows expanded expression of the *D. prolongata Poxn* upAB+ enhancer at the prepupal stage (Fig. 4A); mechanosensory bristles in this region are affected by the *D. prolongata* allele driving *sv* RNAi, but not by the *D. rhopaloa* allele. Other mechanosensory bristles where the *D. prolongata* enhancer has a stronger effect than the other species’ alleles are marked by thin black arrows. F. Counts of mechanosensory (top), chemosensory (middle), and total (bottom) t1 bristles that are affected by *sv* RNAi expression driven by *Poxn* enhancers from *D. prolongata* (red), *D. rhopaloa* (blue), and *D. melanogaster* (green). Sample sizes n_sex,region,species_: n_mAB+p_=29, n_mAB+r_=17, n_mAB+m_=17, n_mB+p_=27, n_mB+r_=11, n_mB+m_=25, n_mAp_=6, n_mAr_=5, n_mAm_=8; n_fAB+p_=9, n_fAB+r_=8, n_fAB+m_=8, n_fB+p_=18, n_fB+r_=11, n_fB+m_=31, n_fAp_=5, n_fAr_=7, n_fAm_=7. Asterisks represent significant differences determined by paired t-tests for comparisons between *D. prolongata* and *D. rhopaloa,* after Bonferroni correction for multiple tests; single asterisks are significant at P = 0.05, double asterisks at 0.01, and triple at 0.005. The *D. prolongata* and *D. rhopaloa* enhancer alleles have similar effects on chemosensory bristles. However, the *D. prolongata* upAB+ allele has significantly higher activity in mechanosensory bristles in both sexes, with a stronger difference in males. The *D. prolongata* upB+ enhancer shows significantly higher activity in mechanosensory bristles in males, but not in females. upA enhancer shows similar activity in *D. prolongata* and *D. rhopaloa.* Interestingly, upAB+ enhancers from both *D. prolongata* and *D. rhopaloa* affect a greater number of chemosensory bristles in *D. melanogaster* females compared to the *D. melanogaster* upAB+ enhancer. The differences observed between both species and *D. melanogaster* presumably reflect older evolutionary changes and are not the focus of this study. G. Dorsal side of the t1 segment of *D. melanogaster* male foreleg showing the effects of driving *Poxn* expression using *D. prolongata* (G) or *D. rhopaloa* (G’) *Poxn* upAB+ enhancers. Driving *Poxn* in the domain of the *D. prolongata* enhancer transforms some mechanosensory bristles in the anterior part of the tarsus into chemosensory bristles (red dashed circles); the same bristles are not affected when *Poxn* expression is driven by the *D. rhopaloa* upAB+ allele (black circles). In contrast to male *D. prolongata* (Fig 1C), most bristles in longitudinal rows retain mechanosensory identity (black arrows).

Images were taken under Brightfield illumination using a Leica DM500B microscope with a Leica DC500 camera, and processed in Adobe Illustrator.

## Results

### Recent male-specific expansion of the leg chemosensory system in *D. prolongata*

The male forelegs of *D. prolongata* are greatly enlarged compared to the other leg pairs, the female forelegs, and the male forelegs of related species (Luecke & Kopp, 2019; Singh & Gupta, 1977). The large male forelegs are employed in species-specific courtship behavior, where the male shakes the female’s abdomen with his forelegs prior to copulation (Setoguchi et al., 2014). The change in size is accompanied by a dramatic, male-specific reorganization of leg sensory organs (Figure 1). In *Drosophila melanogaster* the foreleg first tarsus has 7 chemosensory bristles in females and 10-11 in males (Mellert et al., 2012; Tokunaga, 1962); the number and locations of these bristles are stereotypical and largely conserved across most of the genus, including the closest relatives of *D. prolongata* (Figure 1). The remaining external sensory organs on the foreleg are the much more numerous mechanosensory bristles. In contrast, the first tarsus on *D. prolongata* male forelegs has nearly 50 chemosensory bristles (Figure 1), while the female forelegs have the same chaetotaxy as in other *Drosophila* species. The two types of bristles can be easily distinguished by external morphology. Mechanosensory bristles are straight, pointed, and have triangular bracts at their bases, proximal to the bristle socket; chemosensory bristles are thinner, strongly curved, blunt at the tips, and lack bracts (Figure 1).

To investigate the male-specific expansion of the leg chemosensory system in *D. prolongata* in more detail, we focused on the first tarsal segment (t1), which has well-described, stereotypical chaetotaxy. In the forelegs of *D. melanogaster* and most other species, t1 mechanosensory bristles are arranged into ∼8-10 transverse bristle rows (TBRs) on the anterior-ventral surface, and into seven longitudinal rows in the rest of the segment (Tokunaga, 1962). In males of many species, some of the distal TBRs are modified into sex combs (Kopp, 2011). In *D. prolongata* males, all seven longitudinal rows can still be discerned; however, rows 1, 2, 5, and 6 contain chemosensory as well as mechanosensory bristles (Figure 1). Row 1 is an especially clear example, consisting of mostly chemosensory bristles in *D. prolongata* but entirely mechanosensory in the other species. Additional chemosensory bristles are present at ectopic (non-stereotypical) positions between the conserved longitudinal rows, enough even to form partial longitudinal rows that complicate assigning homology between species. Row 5 in particular appears to be split between two rows of chemosensory bristles, leaving it ambiguous which is the true row 5 homolog. In any case the bulk of row 5 has been transformed into chemosensory organs. No chemosensory bristles are observed among TBRs on the ventral side, however the sex comb is greatly reduced (Figure 1). Thus, the increased number of chemosensory bristles in the male forelegs of *D. prolongata* appears to come from two different sources: a homeotic transformation of mechanosensory into chemosensory bristles, and the development of supernumerary chemosensory bristles akin to those induced by ectopic *Pox neuro* expression (Boll & Noll, 2002). This suggests that in *D. prolongata* males, the developmental program that specifies chemosensory bristles has expanded into sensory organ precursors that follow a deeply conserved mechanosensory fate in other species, as well as into supernumerary SOPs that are novel to *D. prolongata*. We decided to focus on two candidate genes that may contribute to this evolutionary change: *Pox neuro* (*Poxn*) and *doublesex* (*dsx*).

### *Poxn* expression is expanded in *D. prolongata* male forelegs

In the external sensory organs of *D. melanogaster*, *Poxn* is necessary and sufficient for chemosensory fate. Sensory organ precursors (SOPs) that express *Poxn* develop into chemosensory bristles, while SOPs that lack *Poxn* expression become mechanosensory bristles by default (Awasaki & Kimura, 1997). We hypothesized that expanded *Poxn* expression in the developing prothoracic leg of male *D. prolongata* may lead to the homeotic transformation of mechanosensory into chemosensory bristles, while female *D. prolongata* and *D. rhopaloa* would have *Poxn* expression similar to *D. melanogaster*. To test this hypothesis, we compared Poxn protein expression in the t1 segment of the prothoracic leg between *D. prolongata and D. rhopaloa* (Figure 2). Leg bristle SOPs in *D. melanogaster* are specified in two waves: chemosensory and a few largest mechanosensory bristles are present by 5 hours after puparium formation (APF), while the remaining mechanosensory bristles are specified later, at late prepupal or early pupal stages (Held, 1990; Held, 2002). At this stage, we therefore expect all chemosensory SOPs, but only a small minority of mechanosensory SOPs, to express sensory organ markers. This stage represents a narrow developmental window between the full eversion of all tarsal segments and the formation of pupal cuticle impervious to antibodies, and thus can be easily recognized by morphology and histology (Tanaka et al., 2011). At the equivalent stages in our focal species (9 hr APF in male *D. prolongata*, 8 hr APF in female *D. prolongata,* and 6 hr APF in both sexes of *D. rhopaloa*), we observe more extensive *Poxn* expression in the forelegs of *D. prolongata* males compared to females and to *D. rhopaloa* males (Figure 2). Double staining against Senseless (Sens), which marks all SOPs soon after their specification (Nolo et al., 2000), shows that Poxn-positive/Sens-positive cells are much more numerous in *D. prolongata* males than in females (Figure 2 A, B), especially in the dorsal t1 where most male-specific chemosensory bristles develop (Figure 1). Most *Sens*-expressing cells at this stage also express Poxn (Figure 2 A, B). Double staining for Cut, which marks SOPs that have acquired external mechanosensory and gustatory sensory organ fate, reveals many Poxn-positive/Cut-negative cells in *D. prolongata* males (Figure 2C), but not in females (Figure 2D) or in *D. rhopaloa* (Figure 2E-F). Early expression of *Sens* is characteristic of chemosensory bristle development (Held, 2002; Jafar-Nejad et al., 2003), *Poxn* acts as an upstream activator of *cut* in some external sensory organs (Vervoort et al., 1995), and ectopic *Poxn* expression can induce ectopic chemosensory bristles in adult legs (Boll & Noll, 2002). The Sens-positive, Poxn-positive, Cut-negative cells presumably represent chemosensory SOPs at an early stage of development, suggesting that the male-specific expansion of the chemosensory system in *D. prolongata* involved spatial and temporal changes in the deployment of a developmental module that includes *Poxn*. This led us to focus on the evolutionary changes in *Poxn* regulation.

**Figure 2:**
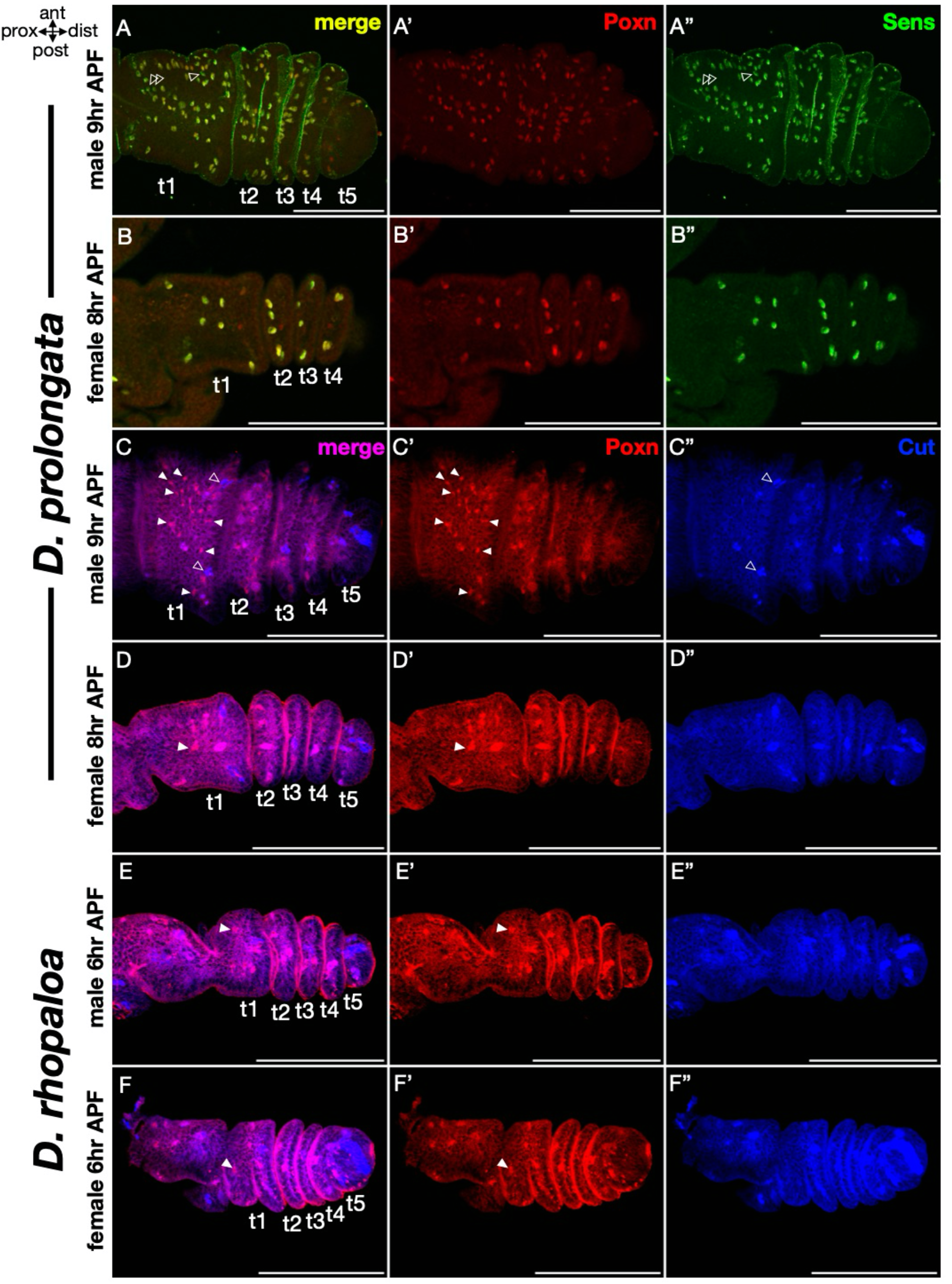
Expression of *Pox neuro* is expanded in the male forelegs of *D. prolongata*. Confocal image stacks of the dorsal surface of pupal prothoracic legs. All legs are oriented with anterior side up; the five tarsal segments are marked t1-t5. All timepoints are equivalent to 5 hours after puparium formation (APF) in *D. melanogaster* (scaled for the total duration of pupal development). Scale bar in all images is 100 micrometers. A-B. Double staining with rabbit Poxn (red) and guinea pig Senseless (Sens) (green). Sens marks all specified SOPs, (Nolo et al., 2000). A. Male *D. prolongata* pupal T1 leg at 9 hr APF. Nearly every cell expressing Poxn also has Sens, indicating that *Poxn* expression in *D. prolongata* is part of a larger sensory organ developmental program. Bristle development occurs over a wide time window, with the basic rule that chemosensory bristles and the largest mechanosensory bristles are specified first (Held 2002). A few nuclei that express Sens but not Poxn (marked with outlined arrowheads) are likely the earliest developing mechanosensory bristles. B. Female *D. prolongata* pupal T1 leg at 8 hr APF. Sens expression is much less extensive than in males, and overlaps Poxn expression entirely. This is consistent with an early pupal stage where only the chemosensory sensory organ precursors (SOPs) have been specified. The sexual dimorphism in both Poxn and Sens expression reflects the lack of chemosensory bristle expansion in females. C-F. Double staining with guinea pig Poxn (red) and mouse Cut (blue). Cut marks SOPs that have acquired external sensory organ fate (Blochlinger et al., 1993); during chemosensory bristle development, Cut expression may appear later than Poxn (Awasaki & Kimura, 1997). A. C. Male *D. prolongata* pupal T1 leg at a slightly earlier developmental stage than in (A). Many but not all Cut-expressing cells express Poxn. Cut-expressing nuclei without Poxn, likely early mechanosensory SOPs, are marked with outlined arrowheads. Poxn expression is also seen in many cells across the t1 segment that do not express Cut (marked with solid arrowheads); the Poxn-positive / Cut-negative cells are likely specified as chemosensory SOPs, but are not as far advanced into the bristle development program as the Cut-expressing cells. B. D. Female *D. prolongata* pupal T1 leg at the same stage as (C). A similar number of Cut-expressing cells are present as in males, with a similar degree of overlap with Poxn expression. Unlike males, only a few Poxn-expressing cells do not express Cut (marked with solid arrowhead), reflecting a smaller number of chemosensory bristles in females. E-F. Poxn and Cut expression in male and female *D. rhopaloa* at 6 hr APF. In both sexes, the overlap between Poxn and Cut expression is similar to female *D. prolongata* (D). The rare Poxn-positive, Cut-negative nuclei are marked with solid arrowheads.

### *Poxn* expression is controlled by non-additive interactions among widely dispersed enhancers

*Poxn* expression in the leg bristles of *D. melanogaster* is controlled by a complex set of *cis*-regulatory elements (CREs) (Boll & Noll, 2002). We first re-examined the *Poxn* locus of *D. melanogaster* using Gal4/UAS transgenic reporters (Figure 3). The boundaries of reporter constructs were placed in regions of high sequence conservation, so that clear homology could be established between *D. melanogaster, D. prolongata,* and *D. rhopaloa* CREs. To reduce the risk of splitting regulatory regions, we extended reporter fragments beyond previously identified CREs and included ample overlap between regions. We restricted our search to the intergenic regions between *Poxn* and neighboring genes. Although this region has been shown to rescue wild type *Poxn* expression in *D. melanogaster* (Boll & Noll, 2002), we cannot rule out the existence of additional, functionally redundant enhancers outside this region.

The upstream region of *Poxn*, upAB+ (Figure 3A’), drives strong expression in Poxn-positive leg bristle precursors, as well as ectopic expression in surrounding epithelial cells (Figure 3C, Figure S1A). This is consistent with the ability of a smaller “over-rescue” region (Figure 3A) to both restore chemosensory bristle development and induce ectopic chemosensory bristles in loss-of-function *Poxn* mutants (Boll & Noll, 2002). The expression pattern of upAB+ in dorsal t1 consists of six clusters, each consisting of one or several SOPs and the surrounding epithelial cells. We refer to these clusters as anterior-proximal (AP), anterior-distal (AD), anterior-central (AC), posterior-central (PC), posterior-proximal (PP), and posterior-distal (PD) (Figure 3C’’).

Comparing the activity of upAB+ (Figure 3C, Figure S1A) with its smaller overlapping sub-fragments (Figure 3D-E, Figure S1B-C, Figure S2) reveals non-additive interactions across this region. The proximal upB+ fragment drives expression in isolated SOPs in dorsal t1 corresponding to the six clusters seen in upAB+, as well as broad expression in the more distal t2 and t4 (Figure 3E, Figure S1C) that is reduced with upAB+ (Figure 3C, Figure S1A). The more distal upA fragment drives expression in broad anterior and posterior stripes of epithelial cells in the dorsal t1 and t2 that encompass the chemosensory bristle clusters (Figure 3D, Figure S1B). Thus, the upAB+ expression pattern is shaped by a combination of positive and negative regulatory elements: sequences in the proximal part of upB+ suppress most but not all of the epithelial expression driven by upA, while sequences in upA suppress the broad distal tarsus expression driven by upB+.

We find that upB+ recapitulates most pupal leg expression clusters in t1 (Figure 3E, Figure S1C) and that the *D. melanogaster* upB+ and upAB+ regions produce similar levels of expression in t1 chemosensory bristles (Figure 4F, *D. melanogaster* data). However other studies have found that driving *Poxn* expression with the upB+ region alone does not fully restore chemosensory bristle development in the legs of *Poxn* mutants (Boll & Noll, 2002) (summarized in Figure 3A), suggesting sequences present in distal upA are necessary for full *Poxn* expression in the leg. For this reason, and because we were unable to identify any smaller fragments that had chemosensory bristle activity without broad ectopic expression, we treat the full upAB+ region as conservative boundaries that contain multiple regulatory elements which drive leg chemosensory bristle expression. We therefore sought to compare the expression produced by the *D. prolongata* and *D. rhopaloa* sequences for the full upAB+ region, as well as the smaller upA and upB+ fragments.

### The upstream *Poxn* enhancers of *D. prolongata* drive expanded gene expression

The upstream *Poxn* region including upAB+ shows well conserved synteny between *D. prolongata*, *D. rhopaloa*, and *D. melanogaster* (Figure 3B), allowing us to make homologous reporter constructs and compare their activity in the common *trans*-regulatory background of *D. melanogaster*. Any differences between *D. prolongata* and *D. rhopaloa* reporters in *D. melanogaster* can be attributed to *cis*-regulatory divergence between these two species. However, given the potential divergence of *trans*-regulatory landscapes between these species and *D. melanogaster*, the differences between *D. prolongata* and *D. rhopaloa* CREs in transgenic *D. melanogaster* may not fully reflect their functional divergence in their native genomic backgrounds. Reporter assays therefore provide a conservative estimate of the contribution of *cis*-regulatory divergence to the evolution of *Poxn* expression.

One possible source of a novel expression pattern could be the origin of a new CRE. To explore this possibility, we generated reporters spanning the entire *D. prolongata Poxn* locus including intronic, upstream, and downstream non-coding sequences (Figure 3A” and Figure S2A). The region immediately upstream of the transcription start site (upCb) and both intronic regions gave no expression in the developing prothoracic leg (Figure S2F-H). The downstream region (downA) drives leg bristle expression that is stronger in males than in females (Figure S2C). When we tested the homologous region from *D. melanogaster*; its expression was similar to the *D. prolongata* reporter (Figure S2B), suggesting that this region includes a conserved bristle CRE. This is consistent with a previous report that sequences downstream of *Poxn* may contribute to its expression in leg bristles (Boll & Noll, 2002). Since the downstream CRE does not show novel or expanded activity in *D. prolongata* males (in fact, it appears to be stronger in *D. melanogaster* (Figure S2B, C)), we continued to focus on the upstream regulatory regions.

In prepupal legs, the upAB+ enhancers from *D. prolongata* and *D. rhopaloa* drive t1 expression that is generally similar to the homologous *D. melanogaster* region, with a few notable differences. The AP, AD, PP, and PD clusters are conserved, while the AC and PC clusters are reduced compared to *D. melanogaster* (Figure 3C, Figure 4A-D). A key difference between the *D. prolongata* and *D. rhopaloa* enhancers can be seen in the AP cluster: in the *D. prolongata* allele, this cluster extends distally into the space between the AP and AD clusters (Figure 4A, white arrowhead), while the *D. rhopaloa* reporter marks an AP cluster similar to the *D. melanogaster* allele (see Figure 3C) and maintains a wide gap between the AP and AD clusters (Figure 4C, grey arrowhead). This difference is only observed in male legs (Figure 4A, C); in females, both *D. prolongata* and *D. rhopaloa* CREs show a wide gap between the AP and AD clusters (Figure 4B, D), similar to males that carry the *D. rhopaloa* reporter (Figure 4C). This *cis*-regulatory difference between *D. rhopaloa* and *D. prolongata* is therefore sex-specific, and could contribute to the increased number of chemosensory bristles in *D. prolongata* males. Moreover, the AP and AD clusters roughly correspond to longitudinal bristle rows 1 and 2, which in *D. prolongata* have the highest numbers of chemosensory bristles in place of the ancestral mechanosensory ones (Figure 1).

To identify the adult bristles that correspond to the clusters of *Poxn* CRE activity in prepupal legs, we used the *Poxn*-Gal4 reporters to drive a UAS-RNAi construct targeting the *shaven* (*sv*) gene, which is essential for bristle shaft development (Kavaler et al., 1999). *sv* RNAi eliminates or truncates bristle shafts, allowing us to link prepupal CRE expression to the adult bristle pattern. None of the reporters consistently affect all chemosensory bristles, although each chemosensory bristle is impacted in at least some individuals (Figure 4E). This is consistent with the earlier report that additional *Poxn* regions are required for robust leg bristle specification (Boll & Noll, 2002), and with the presence of the downstream *Poxn* enhancer (Figure S2B, C). upAB+ enhancers from *D. prolongata* and *D. rhopaloa* affect similar numbers of chemosensory bristles in *D. melanogaster*, in both male and female legs (Figure 4F). However, the *D. prolongata* CRE impacts significantly more mechanosensory bristles compared to the *D. rhopaloa* and *D. melanogaster* alleles (Figure 4F). This difference is especially pronounced on the anterior side of t1 (Figure 4E), corresponding to the merging of the AP and AD expression clusters in prepupal legs (Figure 4A) and to the region where many chemosensory bristles develop in *D. prolongata* males (Figure 1).

As an additional assay of their activity, we used the upAB+ enhancers from *D. prolongata* and *D. rhopaloa* to drive UAS-*Poxn* expression in *D. melanogaster* male forelegs. Using the *D. prolongata* CRE transformed some anterior-dorsal mechanosensory bristles to chemosensory fate (Figure 4G), while the *D. rhopaloa* CRE failed to modify these bristles (Figure 4G’). Together, the *sv* RNAi and UAS-*Poxn* experiments show that functional divergence of the upAB+ *cis*-regulatory region between *D. prolongata* and *D. rhopaloa* may contribute to chemosensory system expansion in *D. prolongata* via a homeotic transformation of mechanosensory into chemosensory bristles.

We then used the *sv* RNAi assay to compare the activity of the upA and upB+ sub-fragments between species. Similar to upAB+, upB+ reporters did not show significant interspecific differences in their effects on chemosensory bristles, but the *D. prolongata* upB+ CRE impacted more mechanosensory bristles compared to the *D. rhopaloa* and *D. melanogaster* alleles (Figure 4F). Consistent with this result the *D. prolongata* upB+ reporter shows somewhat broader expression in male prepupal t1 compared to the *D. rhopaloa* allele (Figure S3C, D). upA reporters from all three species have a similarly strong effect on chemosensory bristles (Figure 4F), consistent with the broad prepupal expression driven by this enhancer (Figure 3D, Figure S3A, B). Surprisingly, the upA reporters from *D. prolongata* and *D. rhopaloa* affect fewer mechanosensory bristles in *D. melanogaster* males compared to the *D. melanogaster* allele, despite the consistently lesser activity of the larger upAB+ CRE in *D. melanogaster* compared to the other species (Figure 4F). This suggests the expanded expression of the upAB+ CRE in *D. prolongata* masks a more complex series of evolutionary changes. An increase in the activity of the upB+ region in *D. prolongata* after its divergence from *D. rhopaloa,* which expanded *Poxn* expression into some ancestrally mechanosensory bristles, may be partly counteracted by the lower “boosting” activity of the upA region in *D. prolongata* and *D. rhopaloa* compared to *D. melanogaster*.

The differences in the expression of the *D. prolongata* and *D. rhopaloa* upAB+ CREs in *D. melanogaster*, while significant (Figure 4), do not fully account for the differences in Poxn protein expression between *D. prolongata* and *D. rhopaloa*, or between *D. prolongata* males and females (Figure 2). The downstream Poxn enhancer cannot explain this disparity (Figure S2B, C). Although *D. prolongata* upAB+ expression in *D. melanogaster* males pales in comparison to Poxn expression in *D. prolongata* males, this CRE nevertheless shows male-specific upregulation (Figure 4A, B). These results indicate that additional loci contribute to chemosensory system expansion in *D. prolongata,* and that at least some of these loci act in a sex-specific manner. We therefore tested whether another candidate gene, *doublesex* (*dsx*), contributes to this evolutionary change.

### Expanded *dsx* expression in *D. prolongata* is due to *cis*-regulatory evolution

The *Drosophila dsx* gene encodes a transcription factor that specifies external morphological differences between males and females via alternatively spliced, male- and female-specific isoforms (*dsxM* and *dsxF*, respectively) (reviewed in (Christiansen et al., 2002)). In the male foreleg, *dsxM* is responsible for converting the distal TBRs into a sex comb (Kopp, 2011; Tanaka et al., 2011) and for specifying male-specific chemosensory bristles (Mellert et al., 2012). This suggests that *dsx* could be involved in regulating sexually dimorphic *Poxn* expression and contributing to the male-specific increase in the number of chemosensory bristles in *D. prolongata*; this regulatory relationship could be indirect, especially since DSX^M^ typically acts as a transcriptional repressor (Christiansen et al., 2002). We find that in prepupal legs, *dsx* shows broader and more diffuse expression in *D. prolongata* males compared to *D. rhopaloa* and *D. melanogaster*, with an especially pronounced increase in SOPs on the dorsal side of the tarsus (Figure 5A-C). Double staining against Sens indicates that *dsx* expression in *D. prolongata* encompasses both SOPs and epithelial cells (Figure 5A), consistent with a potential role for DsxM in activating *Poxn* expression in chemosensory bristles. The broad epithelial expression is also unsurprising given the extreme sexual dimorphism in leg size and pigmentation in this species (Figure 1).

**Figure 5:**
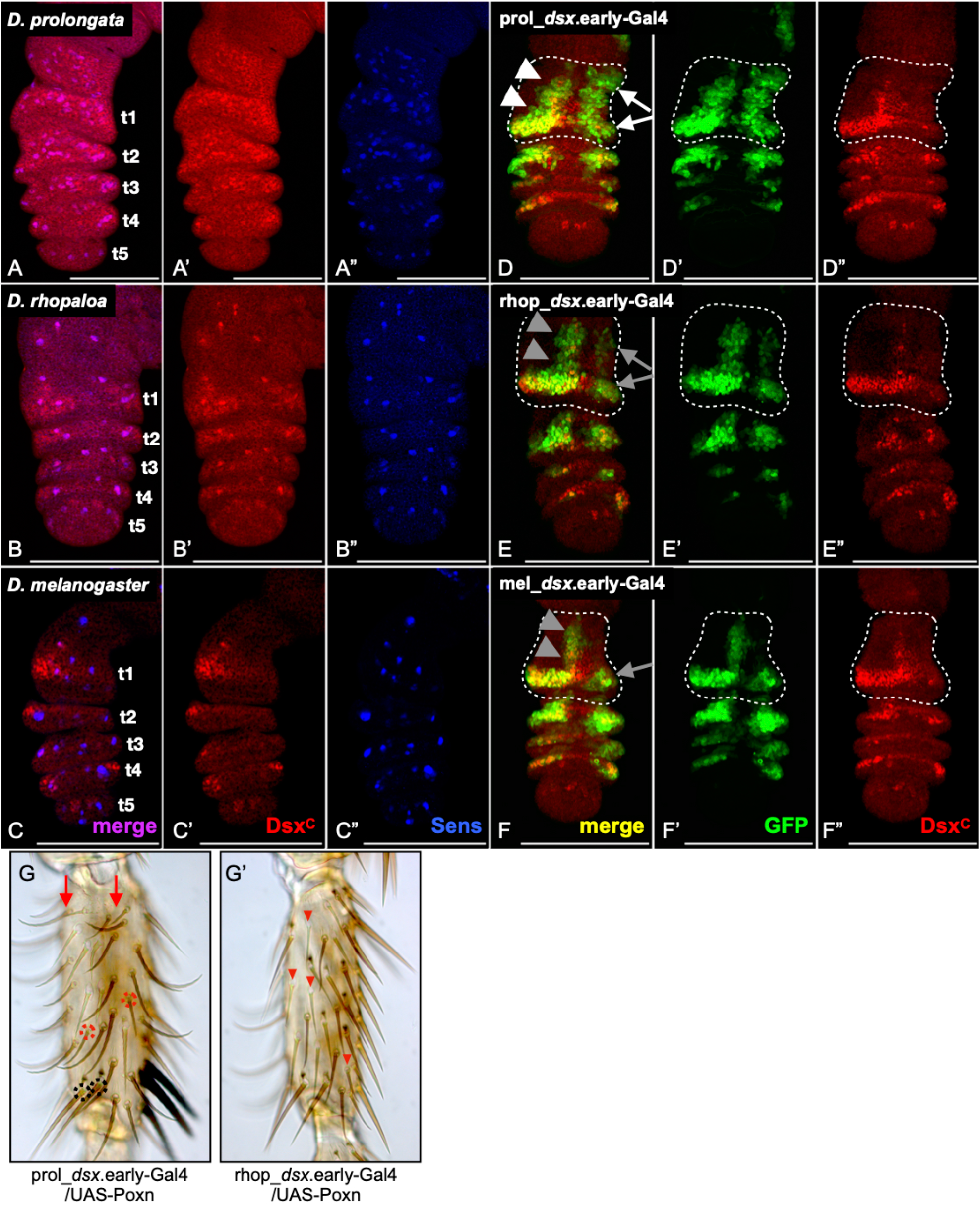
*cis*-regulatory changes contribute to expanded *dsx* expression in *D. prolongata* males. A-C. Single confocal slices of dorsal male forelegs of *D. prolongata* at 9 hr APF (A), *D. rhopaloa* at 6 hr APF (B), and *D. melanogaster* at 5 hr APF (C), representing equivalent developmental stages. Dsx (stained with antibody against the common protein domain, DsxC) is in red and Sens in blue. *D. prolongata* has many more SOPs at this stage, due to the earlier specification of chemosensory bristles (Figure 2). All of these SOPs express Dsx (A, A’). *D. rhopaloa* and *D. melanogaster* have fewer Sens-positive, Dsx-positive cells (B, C). *D. prolongata* males also have broader epithelial Dsx expression (A, A’) compared to the other species (B, C), so SOP Dsx expression stands out less in this species. D-F. Confocal stacks of dorsal male forelegs of *D. melanogaster* at 5 hr APF, with GFP (green) driven by the *dsx* early enhancer from *D. prolongata* (D), *D. rhopaloa* (E), and *D. melanogaster* (F), counter-stained for Dsx protein (red). Enhancers from all three species drive strong expression along the anterior side of the t1 segment (arrowheads in D-F). The *D. prolongata* and *D. rhopaloa* enhancers (D’, E’) show broader anterior expression compared to the native Dsx expression in *D. melanogaster* (D’’, E’’) and to the *D. melanogaster* enhancer (F’). The posterior expression domain of the *D. prolongata* enhancer (white arrows in D) is stronger compared to the *D. rhopaloa* and *D. melanogaster* enhancers (grey arrows in E, F), and extends further proximally than Dsx expression in *D. melanogaster* (D’’). G. Dorsal side of the t1 segment of *D. melanogaster* male foreleg showing the effects of driving *Poxn* expression using *D. prolongata* or *D. rhopaloa dsx* early leg enhancers. Driving *Poxn* in the domain of the *D. prolongata dsx* enhancer (G) produces a nearly complete phenocopy of *D. prolongata* male chaetotaxy (Fig 1C) by converting some longitudinal rows from mechanosensory to chemosensory fate (red arrows), inducing ectopic chemosensory bristles between longitudinal rows (red dashed circles), and reducing the sex comb from 10-12 bristles to 2-3. Some mechanosensory bristles remain at the distal end of t1 (dashed black circles), similar to what is seen in *D. prolongata* males (Fig 1C). Driving *Poxn* with the *D. rhopaloa dsx* early enhancer (G’) disrupts the longitudinal and transverse rows and removes the sex comb entirely. Some ectopic chemosensory bristles are present (examples marked with red arrowheads), but they are not as numerous or organized as those induced by the *D. prolongata dsx* enhancer. The resulting pattern does not resemble the *D. prolongata* (Fig 1C) or *D. rhopaloa* (Fig 1B) bristle phenotypes.

We tested whether, similar to *Poxn*, *cis*-regulatory divergence contributes to the differences in *dsx* expression between *D. prolongata* and *D. rhopaloa.* Our work in *D. melanogaster* has shown that *dsx* expression in the foreleg is controlled by multiple CREs with distinct spatial and temporal activities, including a CRE that acts at the prepupal stage and contributes to both sex comb and chemosensory organ development (G. R. Rice et al., 2019). We compared the activity of homologous CREs from *D. prolongata* and *D. rhopaloa* using GAL4/UAS reporters in transgenic *D. melanogaster.* We find the *D. prolongata* CRE drives much broader expression than the homologous enhancer from *D. melanogaster* (Figure 5D,F; Figure S4D,F). Increased expression is seen on both the anterior and the posterior sides of t1 in both sexes (Figure 5D, Figure S4D). The anterior expression (white arrowheads in Figure 5D) is a slight expansion of a conserved band of *dsx* expression along the ventral/anterior surface of t1 (grey arrowheads in Figure 5F). On the posterior side, the difference is more dramatic: the *D. melanogaster* enhancer is only active in the most distal part of t1 (grey arrow in Figure 5F), while the *D. prolongata* allele drives strong expression more proximally (white arrows in Figure 5D). The broader activity of the *D. prolongata* CRE explains much of the divergence in Dsx expression between these species, and may contribute to the differences in the *trans*-regulatory environment experienced by *Poxn* CREs in *D. prolongata* vs *D. melanogaster*.

The *D. rhopaloa dsx* leg CRE shows broader expression than *D. melanogaster,* but is noticeably weaker than the *D. prolongata* allele, especially in the posterior domain (Figure 5E, Figure S4E). To compare the *D. rhopaloa* and *D. prolongata dsx* enhancers directly, we used them to drive UAS-*Poxn* expression in male forelegs. When the *D. prolongata* CRE was used, a large number of mechanosensory bristles in the t1 segment were transformed into chemosensory bristles (Figure 5G), while the *D. rhopaloa* enhancer produced a much weaker transformation (Figure 5G’). This confirms that *dsx* expression has expanded in *D. prolongata* relative to *D. rhopaloa* due in large part to *cis*-regulatory changes, and shows the tissue experiencing novel expression includes the bristles that have transformed from mechanosensory to chemosensory in *D. prolongata* males.

### Expanded activity of *D. prolongata Poxn* upAB+ CRE is limited to Dsx-expressing cells

Coexpression of *Poxn* and *dsx* in the chemosensory bristles of male *D. prolongata* suggests that *dsx* could be one of the upstream regulators of *Poxn*. To test this hypothesis, we examined UAS-GFP expression driven by the *Poxn* upAB+ CRE from *D. prolongata* and *D. rhopaloa* in *D. melanogaster* male prepupal legs, counterstaining them for the DsxM protein (Figure 6A-D). As described above, the *D. prolongata* upAB+ allele drives epithelial expression between the AP and AD clusters, which is absent in the *D. rhopaloa* allele (arrow in Figure 6A’ vs 6B’). The epithelial cells where the differences between the *D. prolongata* and *D. rhopaloa Poxn* enhancers are observed express DsxM (Figure 6A, B). On the posterior side of t1, the *D. prolongata Poxn* upAB+ CRE drives expression in two SOPs in the PD cluster (Figure 6C), while the *D. rhopaloa* allele shows activity in only one of these SOPs (Figure 6D). The SOP with the differential expression also shows strong DsxM expression (arrowhead in Figure 6C, D).

**Figure 6:**
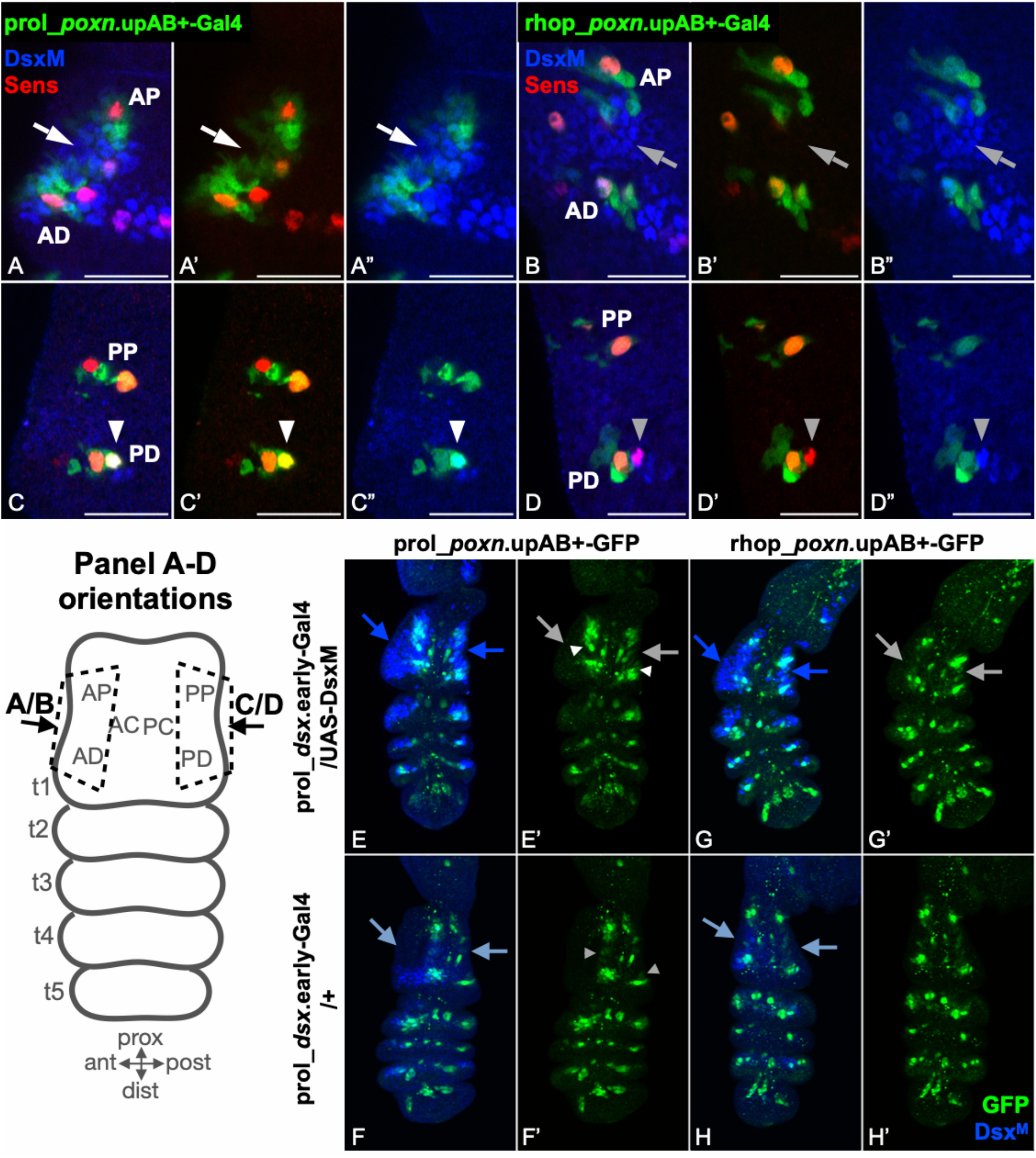
*Poxn* enhancer activity has diverged in Dsx-expressing cells, but expansion of DsxM is not sufficient to expand *Poxn* enhancer activity. A-D. Single confocal slices of *D. melanogaster* male forelegs at 5 hr APF. Poxn_upAB+- Gal4>UAS-GFP.nls is in green, Sens in red, and the male Dsx isoform (DsxM) in blue. Scale bars are 20 micrometers. Orientations are depicted in prepupal leg diagram. A, B. The ventral/anterior surface of the t1 segment showing the AP and AD clusters of upAB+ reporter expression (orientation shown by accompanying diagram). The *D. prolongata* allele (A) drives expression in epithelial cells between these two clusters (arrow), which is absent in the *D. rhopaloa* allele (best seen in A’ vs B’). These cells express Dsx (arrows in A” vs B”). C, D. The dorsal/posterior surface of the t1 segment, showing the PP and PD clusters of upAB+ expression (orientation shown by accompanying diagram). The PD cluster of the *D. prolongata* enhancer covers two SOPs (C), while the *D. rhopaloa* PD cluster covers only one SOP (D). The SOP with differential expression (arrowhead) also shows strong DsxM expression (D’’). E-G. Expansion of DsxM expression is not sufficient to expand the spatial activity of the *D. prolongata Poxn* upAB+ enhancer, but may increase the level of its expression. Z-stacks of dorsal T1 leg surfaces of male *D. melanogaster* at 5hr APF, with GFP in green and DsxM protein in blue. E. UAS-*dsxM* / + ; prol_*dsx*.early-Gal4 / prol_*Poxn*.upAB+-GFP. F. CyO / + ; prol_*dsx*.early-Gal4 / prol_*Poxn*.upAB+-GFP. G. UAS-*dsxM* / + ; prol_*dsx*.early-Gal4 / rhop_*Poxn*.upAB+-GFP. H. CyO / + ; prol_*dsx*.early-Gal4 / rhop_*Poxn*.upAB+-GFP. The expansion of DsxM expression under the control of *D. prolongata dsx* enhancer (dark blue arrows in E, G), relative to wild-type DsxM expression (light blue arrows in F, H) does not expand the activity of either the *D. prolongata* (E’ vs F’) or the *D. rhopaloa* (G’ vs H’) *Poxn* upAB+ enhancer into the region of ectopic DsxM expression. Areas indicated with grey arrows have expanded DsxM expression, but do not show increased *Poxn* enhancer activity. However, some of the existing clusters of *D. prolongata Poxn* upAB+ enhancer expression are more intense in the presence of elevated levels of DsxM (arrowheads in E’ vs F’). The *D. rhopaloa Poxn* upAB+ allele does not show this increase (G’ vs H’). This suggests that the *D. prolongata Poxn* upAB+ enhancer is upregulated by DsxM, but the spatial extent of its activity is not set solely by DsxM.

The fact that expanded activity of the *D. prolongata Poxn* enhancer relative to its *D. rhopaloa* homolog is consistently seen in *dsx*-expressing cells suggests that *dsx* may help delimit the species-specific domains of *Poxn* expression. However, the upAB+ CRE is not active in all Dsx-expressing cells, indicating that *dsx* may be necessary but not sufficient for *Poxn* expression. Expressing *dsxM* in *D. melanogaster* male forelegs under the control of either *D. prolongata* or *D. rhopaloa dsx-*GAL4 CREs (data not shown), or using ubiquitous bristle drivers (Atallah et al., 2014; Tanaka et al., 2011) does not transform mechanosensory into chemosensory bristles. In contrast, expressing UAS-*Poxn* using the same *dsx*-GAL4 drivers results in a strong mechanosensory to chemosensory transformation (Figure 5G, G’). This difference confirms that *dsxM* alone cannot induce *Poxn* expression in SOPs. The need for additional *trans*-regulatory factors, whose expression may also differ between *D. prolongata* and *D. melanogaster*, would explain why the expanded expression driven by the *D. prolongata* upAB+ *Poxn* CRE, while consistent and significant, does not fully reproduce Poxn expression in *D. prolongata* males.

### Expanded *dsx* expression is insufficient to activate the *Poxn* upAB+ enhancer

The observations described so far suggest that at least two evolutionary changes have contributed to the dramatic male-specific expansion of the chemosensory system in *D. prolongata* following its divergence from *D. rhopaloa*: an expanded activity of the *Poxn* upAB+ enhancer in *dsx*-expressing cells (Figure 6), and broader *dsx* expression due to changes in the *dsx* leg enhancer (Figure 5). Next, we asked whether, in addition to these *cis*-regulatory changes, there has also been a change in the interaction between *dsx* and *Poxn* (Figure 7). Specifically, we tested whether the *Poxn* CREs of *D. prolongata* and *D. rhopaloa* differ in their response to increased Dsx expression.

**Figure 7:**
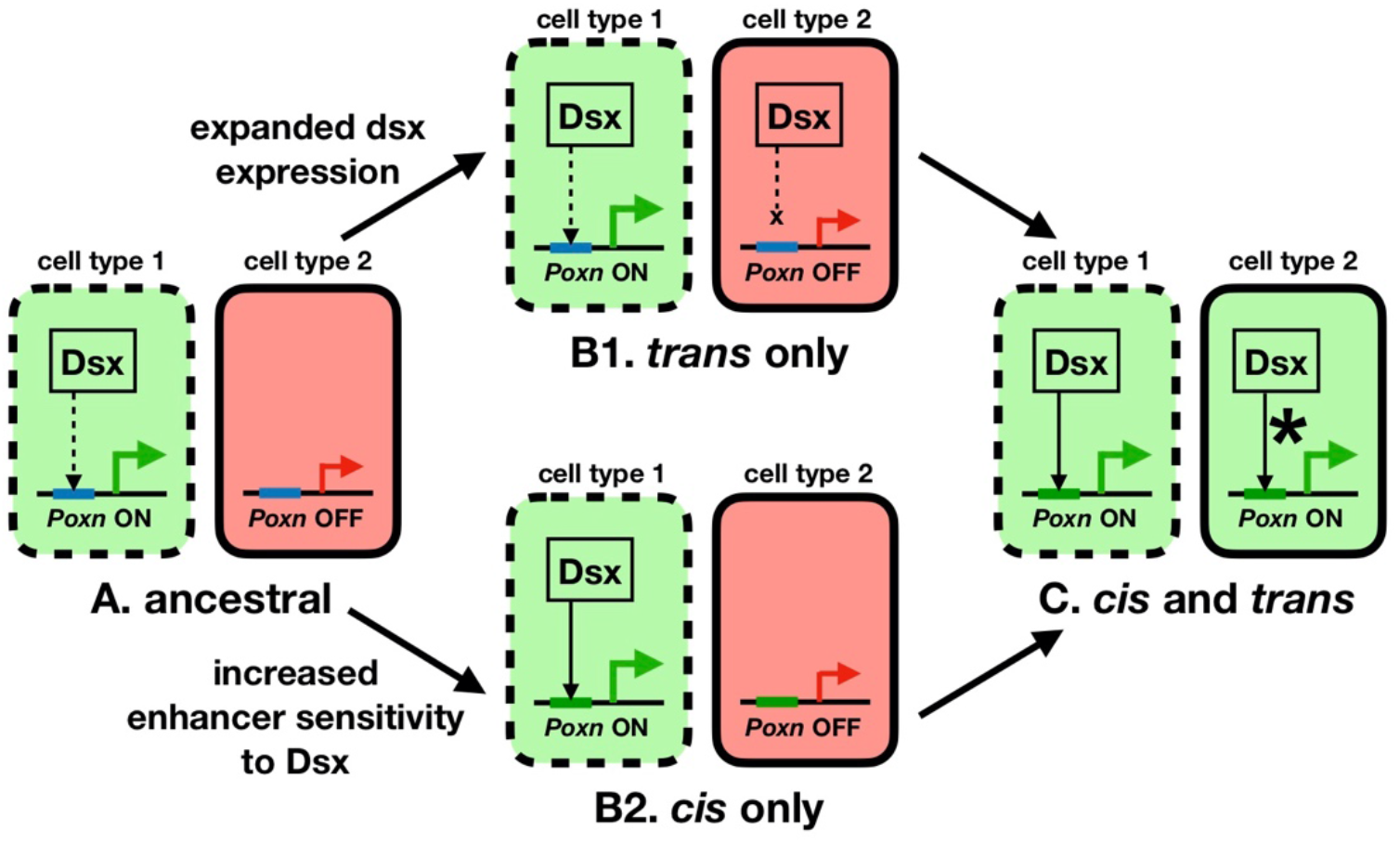
A model of evolving *Poxn* expression through a combination of *cis*- and *trans*-regulatory changes. In the hypothesized ancestral state (A), two cell types have distinct *trans*-regulatory environments, including but not limited to the presence/absence of the DsxM transcription factor. The ancestral *Poxn cis*-regulatory element (blue) activates expression in cell type 1 but not in cell type 2, due in part to positive regulation by DsxM, which may be direct or indirect (dashed arrow). In the course of evolution, expansion of *dsx* expression into cell type 2 does not activate *Poxn* expression in this cell type (B1), because the ancestral *Poxn* enhancer is insensitive to DsxM in the *trans*-regulatory background of cell type 2. The *cis*-regulatory element of *Poxn* may evolve higher sensitivity to DsxM and/or other *trans*-regulatory factors (green enhancer), but this will not result in the activation of *Poxn* expression in cell type 2 due to the absence of DsxM (B2). Expansion of *Poxn* expression into cell type 2 (C) requires a combination of *cis*-regulatory changes (increased sensitivity of the *Poxn* enhancer to DsxM and/or other transcription factors) and *trans*-regulatory changes (spatial expansion of *dsx* expression). Evolution of the *trans*-regulatory landscape must also involve other transcription factors in addition to Dsx (asterisk in C), since experimental manipulation of *dsx* expression alone is not sufficient to expand *Poxn* enhancer activity to the level seen in *D. prolongata* (Figure 6).

We generated transgenic *D. melanogaster* males that carried *Poxn* upAB+ GFP reporters from either *D. prolongata* or *D. rhopaloa*, while also expressing UAS-*dsxM* under the control of the *D. prolongata dsx* enhancer. This manipulation appears to increase the intensity of expression, but is insufficient to expand the expression domains of either the *D. prolongata* or the *D. rhopaloa* upAB+ enhancer (Figure 6). This lack of response has several potential explanations: additional *trans*-regulatory factors may be necessary to activate upAB+; the effect of DsxM on *Poxn* expression may be mediated by a different *Poxn* enhancer; or *dsx* may need to be expressed at the larval stage in order to affect *Poxn* expression in prepupal legs.

## Discussion

Our results suggest that the increased number of chemosensory bristles in *D. prolongata* males results from the expansion of *Poxn* expression, which appears to be produced by interacting *cis*-regulatory changes at *Poxn* and *dsx* along with unknown changes at other loci. Below, we review the evidence in support of this model and discuss how these findings fit into the larger framework of regulatory evolution, and how the organization of CREs can influence their evolutionary potential.

In *D. prolongata*, both *Poxn* and *dsx* expression is expanded into the regions that develop male-specific chemosensory bristles. The CREs of both genes have increased activity in *D. prolongata* compared to *D. rhopaloa*, indicating that *cis*-regulatory changes have contributed to this expansion. However, the difference between *Poxn* enhancer alleles is limited in the *D. melanogaster* background, suggesting that additional changes in *trans*, upstream from *Poxn*, are required for the full expansion of *Poxn* expression in *D. prolongata*. Comparing transgenic reporters in *D. melanogaster* is an inherently conservative test: while all differences in expression can be ascribed to the CRE sequences, these differences may be limited by the divergence of *trans-* regulatory backgrounds between *D. prolongata* and *D. melanogaster*, leading us to underestimate the magnitude of *cis-*regulatory divergence.

We therefore examined candidate *trans-*regulatory gene expression in cells that activate the *D. prolongata* but not the *D. rhopaloa Poxn* enhancer. Intriguingly, the expansion of *Poxn* expression appears constrained by the boundaries of *dsx* expression, suggesting *dsx* is an upstream regulator of *Poxn*. Expressing *Poxn* in the pattern of the *D. prolongata dsx* enhancer produces a phenocopy of the *D. prolongata* chemosensory bristle expansion, consistent with the idea *dsx* acts in *trans* to define the boundaries of *Poxn* expression in *D. prolongata*. However, a *trans*-only model, where Dsx drives expanded *Poxn* expression without any change at the *Poxn* locus, cannot explain our results since ectopic expression of DsxM from the *D. prolongata dsx* enhancer in *D. melanogaster* does not induce ectopic chemosensory bristles. This shows that the *D. melanogaster Poxn* CREs are largely insensitive to increased Dsx expression, although we cannot rule out that increased sensitivity to Dsx evolved before the split between *D. prolongata* and *D. rhopaloa*. Thus, changes in both *cis* and *trans* have contributed to the derived *D. prolongata* phenotype (Figure 7).

We tested this more complex *cis*-and-*trans* model by comparing the activity of the *D. prolongata* and *D. rhopaloa Poxn* enhancers in the background of ectopic DsxM expression. A two-locus additive model predicts increased expression from CRE alleles of both species, while increased expression from only the *D. prolongata* allele favors a model with non-additive interactions. In fact, neither allele responded to increased DsxM expression, indicating that a two-locus *cis*-and-*trans* model is insufficient to explain the evolution of *Poxn* expression. The most likely scenario is that additional upstream regulators of *Poxn*, including perhaps some proneural genes, have also evolved increased expression in *D. prolongata* and are responsible for broader activation of *Poxn* in this species.

Placing our results in a broader context requires a general discussion of enhancer form and function. The two most relevant characteristics of classical enhancers are compactness and tissue-specificity. Compactness refers to the organization of individual enhancers into short stretches of DNA containing clustered binding sites for proteins that interact synergistically to drive transcription independent of other regulatory inputs (Lee et al., 1987; Thanos & Maniatis, 1995). Tissue-specificity describes the ability to recapitulate a specific subset of the gene’s endogenous expression pattern (Geyer & Corces, 1987; Gómez-Skarmeta et al., 1995). These two features combine to produce enhancer modularity, where a single gene can have multiple enhancers that are active in different tissues, and each independent enhancer represents an expression module responsible for a different portion of the gene’s overall expression pattern (Levine, 2010; Long et al., 2016).

Enhancer modularity is a major source of evolutionary potential (Carroll, 2008; Wray, 2007). Many genes are crucial for multiple processes across distinct developmental times and in different tissues. Evolution is highly constrained acting on such genes because changes that are beneficial in some developmental contexts are likely deleterious in others. If, however, these complex expression patterns are governed by modular enhancers, gene expression in any one domain can evolve independently of the others, breaking the pleiotropic constraint. This model is consistent with many examples of morphological evolution driven by divergence in *cis*-regulatory sequences (Jeong et al., 2008; Koshikawa et al., 2015; Prud’homme et al., 2006; Rebeiz et al., 2009), or by gain and loss of entire enhancer activities (Chan et al., 2010; Indjeian et al., 2016; Wallbank et al., 2016).

While these results strongly uphold the evolutionary potential of modular enhancers, recent work has revealed departures from classical enhancer modularity. These deviations fall into four overlapping classes: dispersed enhancers, pleiotropic enhancers, redundant enhancers, and non-additively interacting enhancers. Below, we consider how these deviations might influence regulatory evolution.

Classical enhancers can be remarkably compact. For example, an 813 bp enhancer of the *Drosophila decapentaplegic* (*dpp*) gene faithfully recapitulates *dpp* expression in the embryonic midgut; a 419 bp fragment drives weaker expression in the same spatial; and even a 45 bp sub-fragment largely retains its tissue-specificity (Manak et al., 1995). The “wing-spot” enhancer of the *D. biarmipes yellow* gene can be reduced to 196 bp without loss of spatial pattern (Arnoult et al., 2013). The “eve stripe 2” enhancer, which drives expression of *even-skipped* in one of its seven stripes in the *Drosophila* blastoderm, is 480 bp in length (Small et al., 1992). In contrast, the dispersed “7-stripe” enhancer that drives the blastoderm expression of *runt* in all seven stripes extends over 5 kb, and can neither be split into stripe-specific modules nor truncated without losing spatial accuracy (Klingler et al., 1996). The sex comb enhancer of the *Drosophila doublesex* gene can be reduced to no less than 3 kb (G. R. Rice et al., 2019). This is not a property of particular tissues or spatial patterns – the “zebra element” of the *fushi tarazu* gene, which also drives 7-stripe expression in the *Drosophila* blastoderm, is only 750 bp in length (Dearolf et al., 1989), and the leg enhancer of *bab2*, whose pattern overlaps with the *dsx* sex comb enhancer, is 567 bp (Baanannou et al., 2013). These different degrees of compactness may influence enhancer evolution. If the dense encoding of regulatory information in compact enhancers constrains their evolution and biases the phenotypic outcomes of sequence changes (Fuqua et al., 2020), dispersed enhancers may exhibit faster and less predictable evolution. Another effect of low compactness is the increased chance of overlap with neighboring enhancers that are active in other tissues. To the extent modularity depends on physical separation, this overlap may increase the pleiotropy of an enhancer.

Compact enhancers may still lack modularity if they generate expression in multiple developmental contexts. Such pleiotropic enhancers are increasingly recognized as a relatively common phenomenon (Glassford et al., 2015; Lonfat et al., 2014; Preger-Ben Noon et al., 2018; Rice & Rebeiz, 2019). They can be seen as a natural consequence of the processes underlying developmental evolution in two ways (Rice & Rebeiz, 2019). First, the origin of new enhancers parallels the origin of new genes: while *de novo* origin from ancestrally non-regulatory sequences may occur (Arnold et al., 2014; Villar et al., 2015), most new enhancers arise *via* repurposing of ancestral enhancers by duplication, transposition, or by coopting and expanding the activity of another enhancer (Carroll, 2005; Long et al., 2016; Rebeiz & Tsiantis, 2017). Enhancers that originate in this fashion are likely to retain some of their ancestral functions. Enhancer pleiotropy may also arise during cooption of gene regulatory networks. Expression of a master regulator may induce its downstream developmental module *via* the enhancers of target genes regardless of developmental context (Glassford et al., 2015; Monteiro & Gupta, 2016). Once that module is established in a new tissue, each of these enhancers acquire a second expression domain. Subsequent evolution is then constrained by the pleiotropy of its dual roles. In the most extreme cases, a single transcription factor binding site may affect the activity of a pleiotropic enhancer in multiple tissues (Nagy et al., 2018; Preger-Ben Noon et al., 2018), which would likely impose a strong evolutionary constraint.

Redundant or “shadow” enhancers are also a common feature of gene regulation (Cannavò et al., 2016). The impact of enhancer redundancy on their evolutionary potential has been alternately proposed as a source of potential variation akin to duplicated genes (Hong et al., 2008) or as a barrier to mutational impact on gene expression (Preger-Ben Noon et al., 2016). In one example the loss of an expression domain generated by several distinct and partially redundant enhancers required independent mutations in at least five CREs (Frankel et al., 2011; McGregor et al., 2007). These enhancers act in a largely additive manner such that each mutation influences the phenotype, which presumably gives selection traction to overcome the robustness barrier. Even in cases where enhancers drive apparently identical expression, their combined activity may be essential for spatial precision (Perry et al., 2011) or for robustness to environmental perturbations (Frankel et al., 2010; Perry et al., 2010). Comparison of the two redundant enhancers of *Krüppel* across *Drosophila* species shows that while their combined activity is strongly conserved their separate activities are not (Wunderlich et al., 2015), suggesting that the evolution of these enhancers is constrained by selection rather than by any of their inherent features. Thus, redundant enhancers may imbue gene regulation with an additional layer of flexibility without limiting the rate of regulatory evolution.

The most intriguing deviations from modularity involve non-additive interactions among separate enhancers, in some cases essential for generating correct spatial expression. A clear example of non-additivity is found in the *Drosophila sloppy-paired 1* (*slp1*) gene (Prazak et al., 2010). Segmental stripes of *slp1* expression are driven by two enhancers, proximal and distal, separated by ∼4 kb. The distal enhancer is active in all parasegment stripes but also drives ectopic expression outside of the correct pattern, while the proximal enhancer alone drives reporter expression only in even-numbered parasegments. When combined the proximal enhancer suppresses the ectopic expression, resulting in correct *slp1* expression plainly different from the sum of their separate activities (Prazak et al., 2010). Similarly, *snail* expression in the *Drosophila* embryonic mesoderm is controlled by two non-additively interacting enhancers (Dunipace et al., 2011). In this case, the distal enhancer prevents ectopic spatial expression from the proximal enhancer, while the proximal enhancer dampens the quantitative output of the distal enhancer; only both enhancers acting together generate correct *snail* expression (Bothma et al., 2015; Dunipace et al., 2011).

A common theme emerging from these and other studies of non-additive enhancer interactions is the restriction of enhancer activity by silencing sequences located outside the enhancer boundaries. Transcriptional silencers are common genomic elements, and many regulatory sequences can function as both enhancers and silencers depending on the tissue context (Gisselbrecht et al., 2020). Changes in silencers can be an important avenue for phenotypic evolution. For example, expression of the *ebony* gene, which plays an important role in establishing *Drosophila* color patterns, is controlled by a broadly active enhancer and two well-separated silencers that restrict its spatial activity; all three elements acting together are necessary for correct *ebony* expression (Ordway et al., 2014; Rebeiz et al., 2009). Some intra- and interspecific variation in pigmentation is due to changes in one of these silencer elements, producing broader *ebony* expression (Johnson et al., 2015). Thus, the interaction between activator and silencer elements is central to the mechanism of evolutionary change.

All these departures from classical enhancer modularity can be accommodated under the established theory of phenotypic evolution by focusing on its central claim: of all the mutations that can potentially impact a selected trait, those with the optimal (often but not always lowest) level of pleiotropy are most likely favored (Kopp, 2009; Martin & Orgogozo, 2013). *cis*-regulatory sequences have exceptional evolutionary potential because they are less pleiotropic than coding sequences, not because they lack pleiotropy entirely. Furthermore, enhancers themselves may vary in their evolvability based on their architecture or their pleiotropic functions. This is the perspective from which we revisit our results.

Both *Poxn* and *dsx* expression are expanded in *D. prolongata* males compared to *D. rhopaloa*. Both genes were likely necessary for the evolution of male-specific chemosensory bristles, as *Poxn* sits at the top of the chemosensory bristle developmental module while *dsx* is the main regulator of morphological sexual dimorphism. However, *Poxn* enhancers show only minor differences in activity in a shared (*D. melanogaster*) *trans*-regulatory background, while *dsx* enhancers recapitulate nearly the entire expression difference between the two species. We propose that the regulatory architectures of the two genes may affect their evolutionary malleability. In essence, *Poxn* leg CREs have high mutational potential due to their size, redundancy and non-additive architecture, but also a greater potential for pleiotropic effects due to overlap with other enhancers, whereas the modular *dsx* enhancers present a more compact mutational target but also a more limited potential for pleiotropic consequences. In this instance, it appears that the compact, less pleiotropic enhancer was better able to respond to selection. Tellingly, the changes at *dsx* do not escape pleiotropic effects entirely. The *dsx* early enhancer is active in the developing sex comb as well as chemosensory bristles, and the gain of chemosensory bristles in *D. prolongata* comes at the expense of the sex comb. This tradeoff was evidently tolerated by selection. It is not the pleiotropy *per se* that imposes evolutionary constraint, but the degree to which pleiotropic effects are selectively unfavorable.

The largest caveat to this interpretation is our inability to test the divergence of *Poxn* CREs in the most relevant contexts: the developing legs of *D. prolongata* and *D. rhopaloa*. A truly constrained enhancer would show minimal divergence in all *trans*-regulatory backgrounds. Testing this prediction by developing transgenics in these species would be a valuable addition to our understanding of the rules for enhancer evolution. As they stand at present, our results are consistent with several predictions of evo-devo theory. We show that redeployment of a developmental module underlies the evolution of a complex phenotype, and that *cis*-regulatory evolution of at least two genes has contributed to this redeployment. The remaining questions point to the continued need to explore the mechanisms underlying cooption of developmental modules as a source of phenotypic diversification, as well as the causes and consequences of enhancer pleiotropy.

---

**Supplemental Figure 1:**
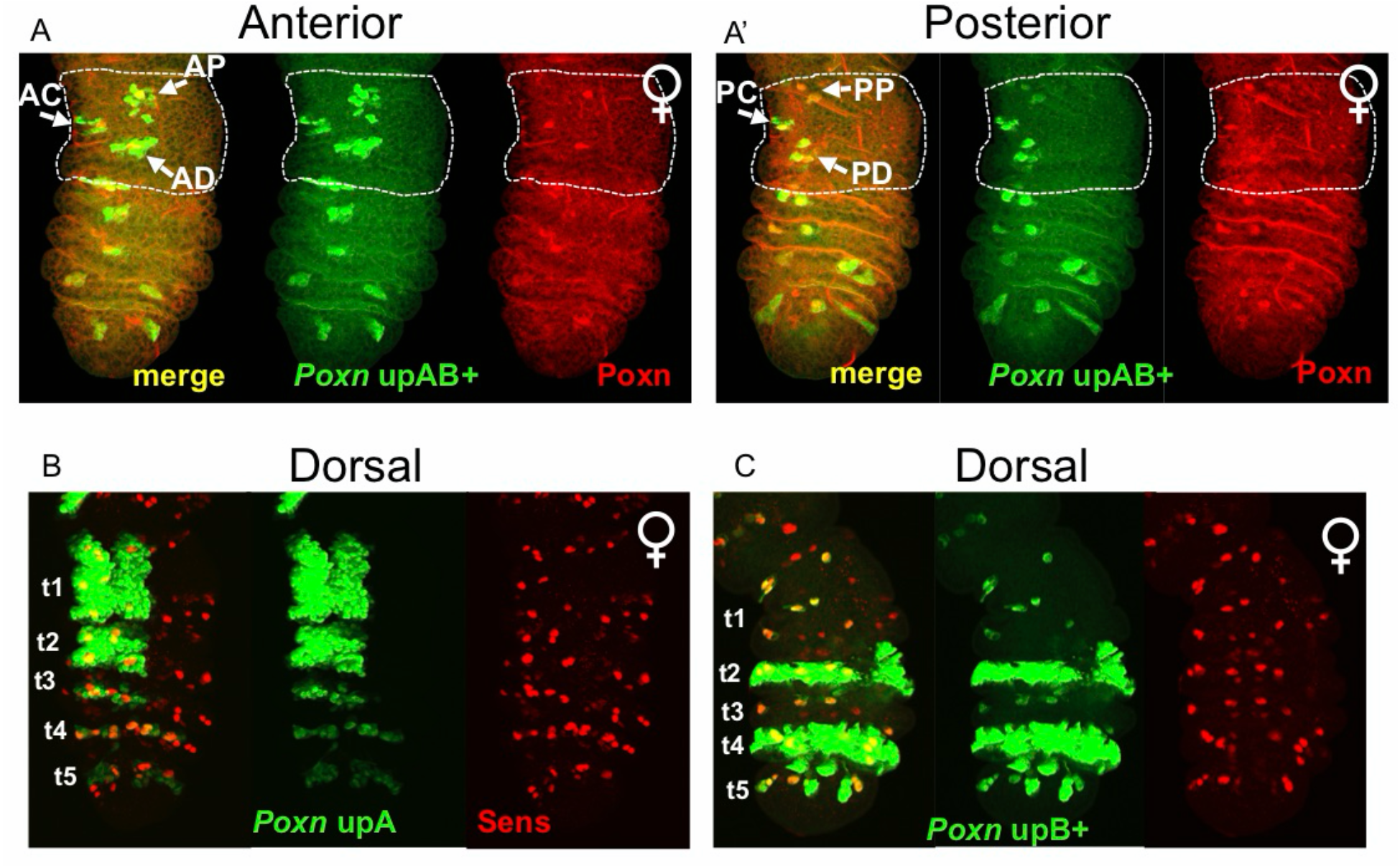
*D. melanogaster Poxn* upAB+ expression in female legs is similar to male expression, but more spatially restricted. A. Z-stacks of the anterior (A) and posterior (A’) *D. melanogaster* female foreleg surfaces at 5 hr APF, showing the activity of the *D. melanogaster Poxn* upAB+ region. Dorsal is left, and t1 segment is circled. *Poxn*-Gal4 > UAS-GFP.nls is in green, and Poxn protein expression in red. Expression clusters are homologous to those in males, and are labeled accordingly. However, the size of the clusters is smaller, especially on the posterior side (compare to Fig. 3C’’). B. Z-stack of the dorsal surface of *D. melanogaster* female foreleg at 5 hr APF, showing the activity of the *D. melanogaster* upA region. *Poxn*-Gal4 > UAS-GFP.nls is in green, and Sens protein in red. C. Z-stack of the dorsal surface of *D. melanogaster* female foreleg at 5 hr APF, showing the activity of the *D. melanogaster* upB+ region. Staining as in (B).

**Supplemental Figure 2:**
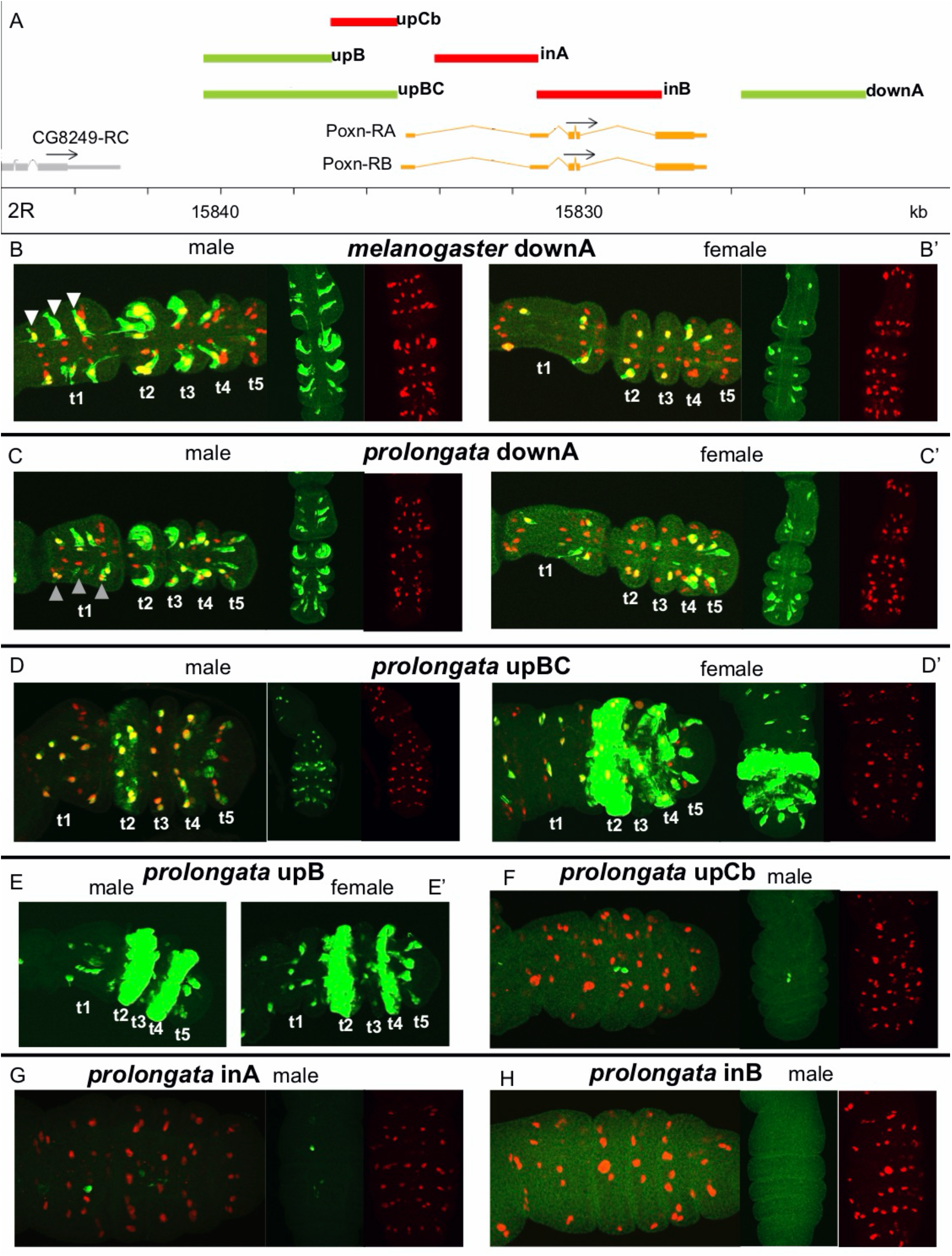
*Poxn* regulatory regions in *D. prolongata* are qualitatively similar to homologous regions in *D. melanogaster*. A. Boundaries of reporter fragments relative to the *Poxn* locus (see also Figure 3A). B-C. Z-stacks of the *D. melanogaster* dorsal T1 leg surface at 5 hr APF showing the activity of *Poxn* downA reporters in both sexes. *Poxn*-Gal4 > UAS-eGFP is in green, Sens protein in red. Anterior is up in the merged images. The *D. melanogaster* downA allele (B) shows expression in and around SOPs that are similar to but not completely overlapping the SOPs marked by the upAB+ enhancer (Figure 3C, E). The downA expression pattern includes several anterior male-specific SOPs in the t1 segment where upAB+ is not active (white arrowheads in B). The *D. prolongata* downA allele also drives expression in these SOPs (C), but at a lower level than the A. *D. melanogaster* allele, particularly on the posterior side (grey arrowheads). Female expression (B’, C’) is less extensive than in males for both species. Sexual dimorphism is greater for the *D. melanogaster* than for the *D. prolongata* enhancer allele, making the downA region an unlikely candidate for increased *Poxn* expression in *D. prolongata* males. B. D. Z-stacks of the dorsal T1 leg surface showing the activity of the *D. prolongata Poxn* upBC enhancer in both sexes. Staining as in B, anterior is up and proximal is left. Expression is similar to that of *D. prolongata* upB (E) and upB+ (Supplemental Figure 3C). C. E. Z-stacks of the dorsal T1 leg surface showing the activity of the *D. prolongata Poxn* upB enhancer in both sexes. Staining as in B, anterior is up and proximal is left. Expression is similar to that of *D. prolongata* upBC (D) and upB+ (Supplemental Figure 3C). F-H. Z-stacks of the dorsal T1 leg surface showing the activity of the *D. prolongata Poxn* upCb, inA, and inB reporters in males. Staining as in B, anterior is up and proximal is left. No SOP expression is seen from any of these enhancers

**Supplemental Figure 3:**
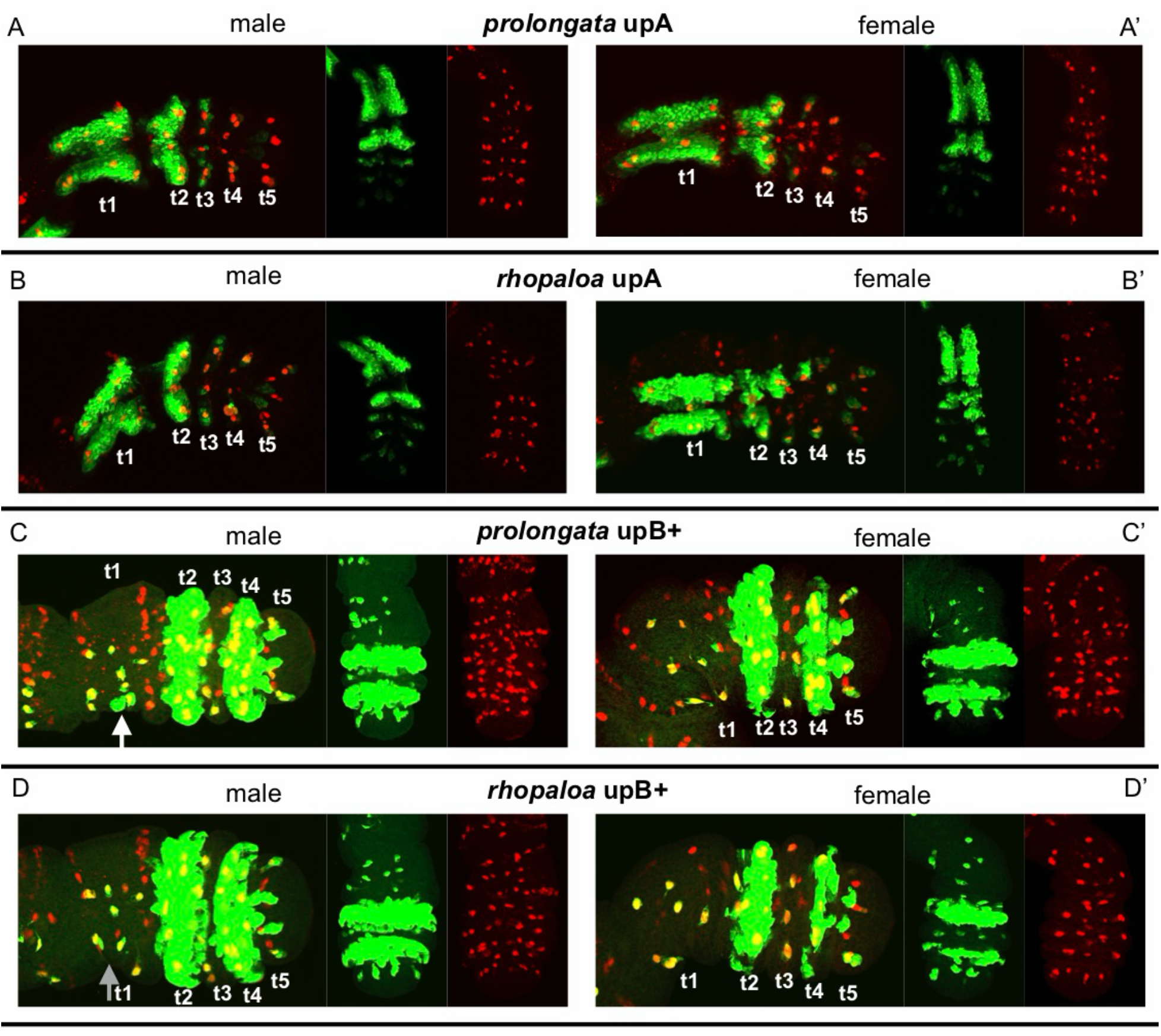
Comparison of *Poxn* enhancers between *D. prolongata* and *D. rhopaloa*. *Poxn*-Gal4 > UAS-eGFP expression is in green, Sens protein in red in all images. Anterior is up, proximal is left in merged images. A. Z-stacks of the *D. melanogaster* dorsal T1 leg surface at 5 hr APF showing the activity of the *D. prolongata Poxn* upA fragment. B. *D. rhopaloa* upA allele. C. *D. prolongata Poxn* upB+ fragment. Expanded expression is seen on the posterior side of the male t1 segment, outside of any specified SOP (white arrow). D. *D. rhopaloa* upB+ allele does not show expression outside posterior t1 SOPs (grey arrow).

**Supplemental Figure 4:**
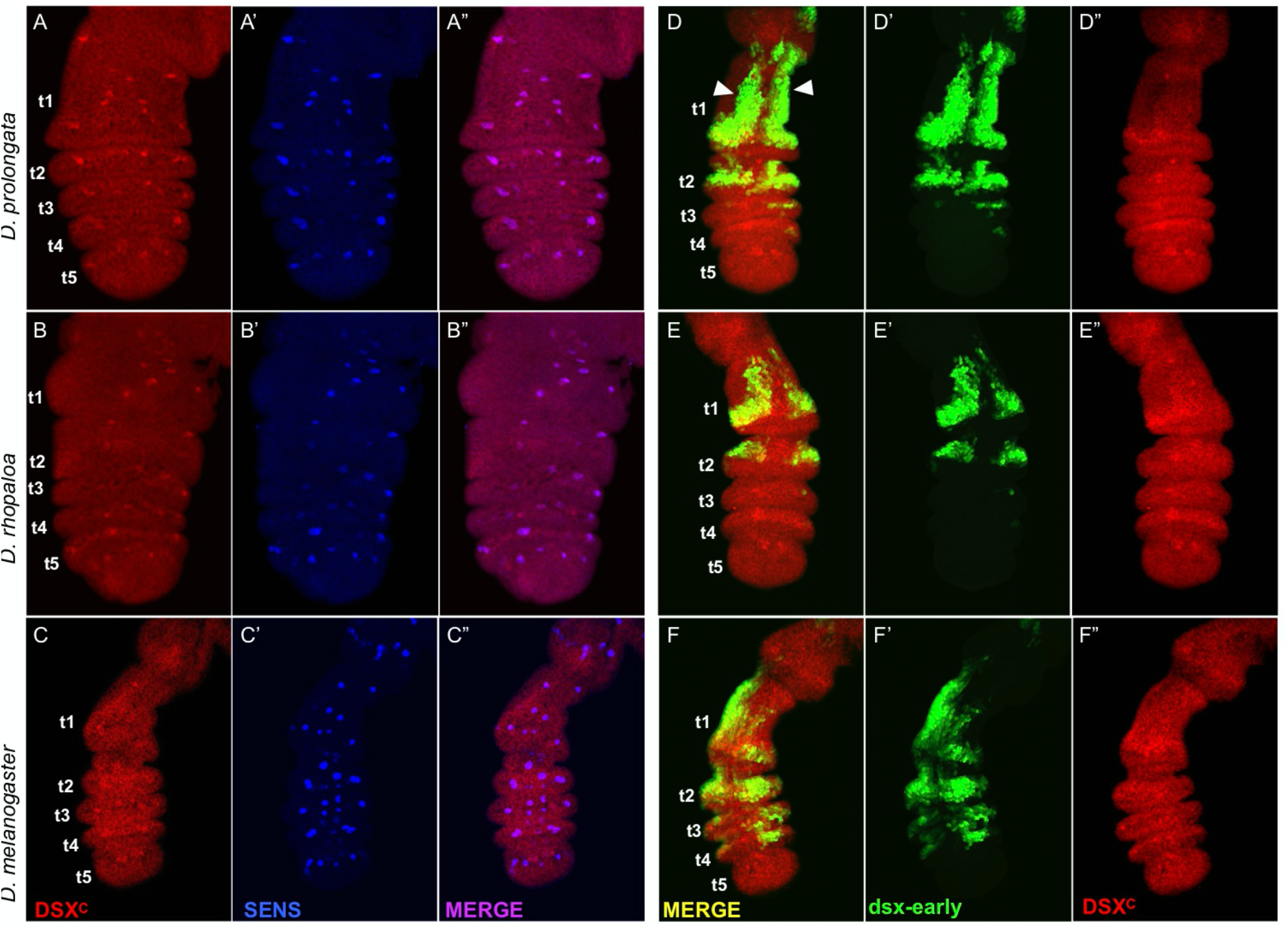
Dsx expression in female *D. prolongata* forelegs and the activity of the *dsx* early foreleg enhancer in females. A-C. Single confocal slices from the dorsal side of female forelegs of *D. prolongata* at 8 hr APF (A), *D. rhopaloa* at 6 hr APF (B), and *D. melanogaster* at 5 hr APF (C, also see D” E”and F” for deeper stacks), stained for DsxC in red and Sens in blue. All ages are equivalent developmental stages. Dsx and Sens co-localize in all species, indicating that these early-specified SOPs have a sexually dimorphic fate. Female *D. prolongata* do not show an expansion in the number of SOPs at this stage (compare to males, Fig. 5A). D-F. Confocal stacks of *D. melanogaster* female dorsal foreleg surface at 5 hr APF. GFP driven by the *dsx* early foreleg enhancer from *D. prolongata* (D), *D. rhopaloa* (E), and *D. melanogaster* (F) is in green, and Dsx protein in red. Expression driven by this enhancer in females is similar to that seen in males (Fig. 5D-F). The *D. prolongata* enhancer drives stronger expression in the proximal t1 (white arrowheads) compared to the other two species.

**Supplemental Table 1:**
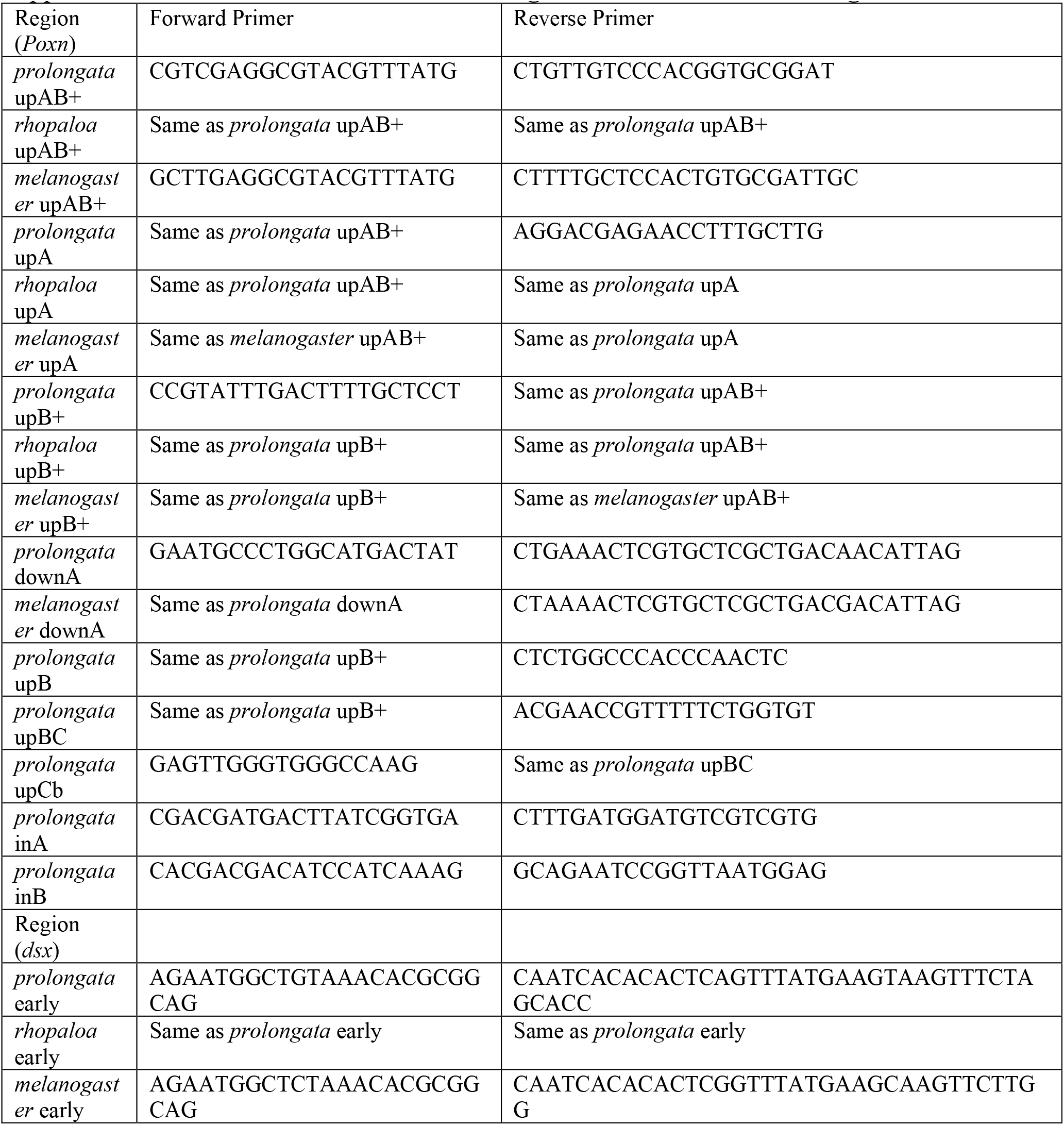
Primers used for cloning *Poxn* and *dsx* enhancer regions.

